# Time-series models can predict long periods of human temporal EEG responses to randomly alternating visual stimuli

**DOI:** 10.1101/2025.08.08.669319

**Authors:** Richard R. Foster, Connor Delaney, Dean J. Krusienski, Cheng Ly

## Abstract

Visual stimuli with constant temporal frequency input is known to induce peaks in the driving frequency of the power spectrum of the electroencephalogram (EEG) over the visual cortex. While EEG responses with random temporal frequencies (m-sequences) have been studied, the underlying mechanisms that shape these responses are not fully understood. We analyze our new EEG data from a controlled experiment with m-sequence inputs and model the EEG using statistical time series models: an autoregressive (AR) model, adding exogenous input to AR (ARX), adding moving average terms (ARMAX), and finally adding a seasonality term (SARMAX). We implement computational methods to robustly handle model instabilities induced by this data, fitting these models with the Box-Jenkins methodology and assessing prediction accuracy for long periods of several seconds out-of-sample. We find in-sample fits are good in all models despite the complexities of the visual pathway, and that all models can predict aspects of EEG: including the distribution of point-wise values in time, the point-wise Pearson’s correlation of EEG and model, and the frequency content. Surprisingly, we find little variation in the performance among these models, with the most sophisticated model (SARMAX) performing comparatively poorly in some instances. Our results suggest the simplest AR model is viable and can out perform more complicated models. Since these models are relatively simple and more transparent than contemporary models with numerous parameters, our study could inform future mechanistic studies of the temporal dynamics of human EEG responses to visual stimuli.

## 1 Introduction

The human visual pathway is a complex system whereby photons of light are transduced in the retina into a series of electrochemical signals relayed to the cortex where the animal ultimately perceives the visual information. While measurement from various neuroimaging modalities can provide insights into the processes occurring in the visual cortex of the brain, scalp EEG has been shown to capture responses to certain visual stimuli reliably, with high temporal resolution. While detailed biophysical models exist for various portions of the visual pathway, no end-to-end biophysical model - from visual input to EEG output - currently exists. In order to better characterize and understand this input-output relationship, EEG were collected from able-bodied participants while they observed simple, stereotyped visual stimuli.

Here we focus on the temporal aspects of the visual stimuli by systematically manipulating the temporal alternating patterns of a fixed spatial stimulus. Participants are instructed to focus on a visual stimulus presented in a head-mounted display to minimize peripheral visual information and distraction. To elicit robust EEG responses, we use a spatially localized checkerboard pattern centered in the visual field that alternates between black and white according to prescribed temporal patterns (Waytowich et al., 2016). In addition to the well-known phenomena of eliciting EEG responses using a fixed alternating frequency (termed steady-state visually evoked potential (**SSVEP**)), we collected EEG data whereby the visual stimuli alternated according to pseudo-random m-sequences using the protocol for code-based visual evoked potentials (**cVEP-BCIs**) in brain-computer interfaces (Bin et al., 2009). Modeling of these EEG responses is less established and could provide insights into the underlying biological systems that process these stimuli.

A natural first step to investigating this complex, multi-scale visual pathway is to use a statistical time series model rather than a more complicated model. Time-series models are ideal for fitting to ample data and can readily be validated by their prediction power of (out-of-sample) data. Although mechanistic models are powerful for testing theories of neural attributes that control/affect measured outputs, a model that faithfully captures the pathway from input frequencies of the visual stimuli to the EEG response remains elusive. To this end, an appealing model is the auto-regressive model (AR) because it is well-known to be the minimally presumptive model (maximum entropy model) that captures the autocorrelation of a single time series (Choi and Cover, 1984; Martini et al., 2024). Although AR models have been used to capture EEG for many years (Birch et al., 1988), the novelty of our framework is in considering complex variants of AR that have components that mimic known mechanics. We use: i) simple auto-regressive (**AR**), ii) adding an exogenous input to model the input stimuli (**ARX**), iii) ARX model with moving-average terms (**ARMAX**) that capture past error corrections, iv) including a seasonality term (**SARMAX**). Components of these models capture several known attributes: neural systems have temporal history dependence (auto-regressive), the exogenous input captures the external stimuli, activity can account for output errors (moving-average), and seasonality captures the periodic nature of the pseudo-random m-sequences that elicit responses at that frequency.

We systematically fit our EEG data to each of the four models and assess their performance in predicting out-of-sample data, only to find little variation in model performance; some of the poorest estimates are generated by the most complicated model SARMAX. Collectively across 15 subjects who each viewed pseudo-random m-sequences for multiple 30-second trials, we had ample data to assess in-sample fits and out-of-sample estimates (300 in total) for each model for a relatively long period of time (7 s in sample, 3.5 s out-of-sample). Model performance was measured via comparisons of model and EEG point-wise values, Pearson’s correlation, and power spectrum (only possible with a relatively long out-of-sample estimate of several seconds). Taken together, our results suggest the AR model is appealing for this data, given its simplicity and that the auto-regressive terms describe how this EEG data is related to its past values.

## 2 Results

EEG data were collected from the visual cortex area while participants viewed a binary (i.e., black and white) visual stimulus for which contrast alternated at fixed and pseudo-random temporal frequencies, i.e., m-sequences (Golomb and Gong, 2005). Participants donned a head-mounted display, where the stimulus was presented in the foveal receptive field (Fig. 1A–B). To analyze the m-sequence responses, EEG signals were spatially filtered via canonical correlation analysis (CCA) against the average EEG response to obtain a set of optimal canonical weights for each electrode (Waytowich and Krusienski, 2015). Linearly combining the 8 individual electrode recordings with the canonical weights gives a single EEG time series that is maximally correlated with the average EEG response (Fig 1C). For the rest of this paper, we use the CCA-spatially filtered EEG when referring to EEG responses, focusing on a single time series. The CCA-spatial filtering is known to provide good signal-to-noise (Johnson and Krusienski, 2018).

**Figure 1.**
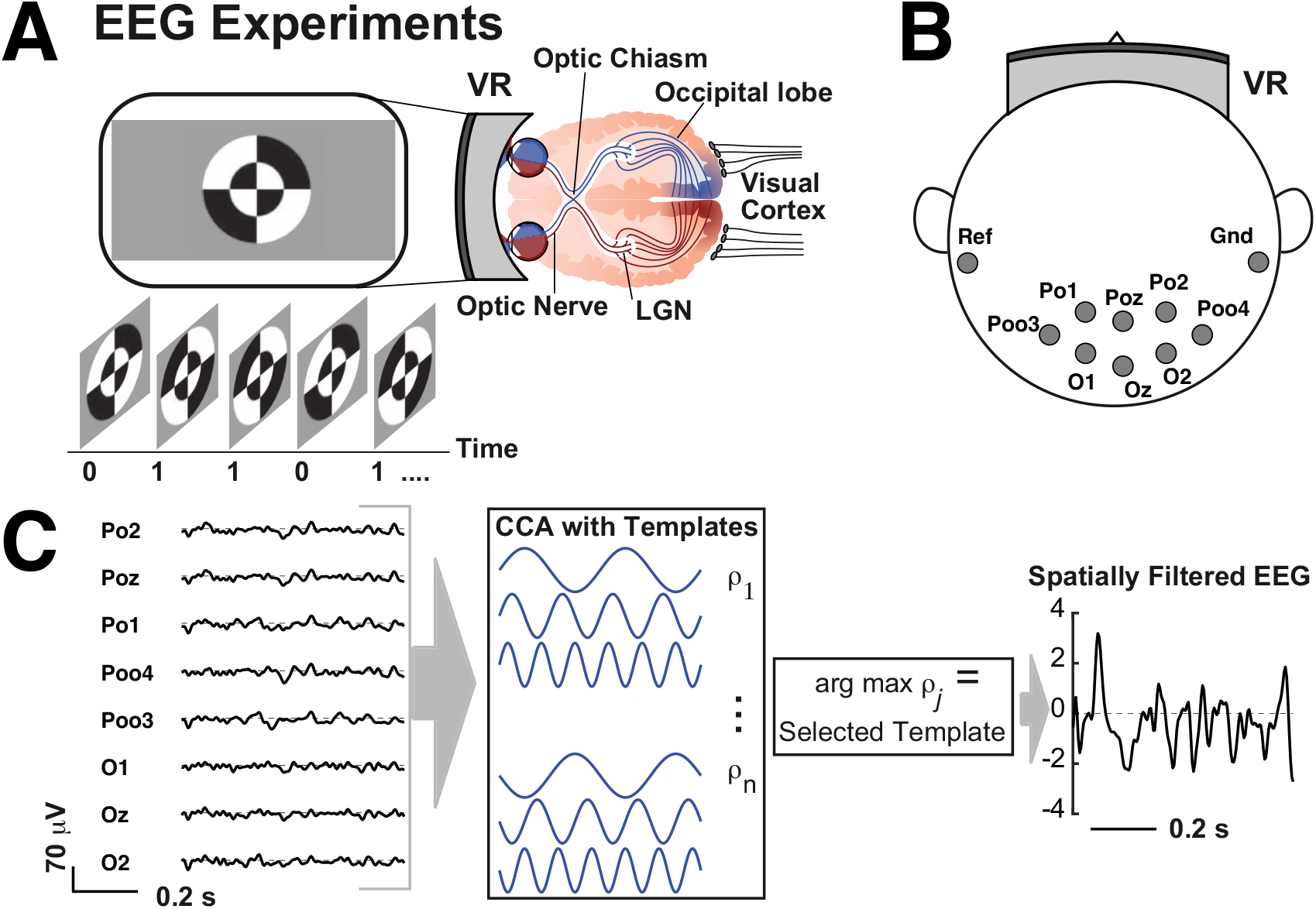
Experimental setup to collect EEG data. A) Participants don a head-mounted display and focus on concentric circle stimuli in the middle of their foveal receptive fields that oscillate between 2 distinct images (black → white, white → black) with rich temporal frequencies – the 0 and 1 labels underneath merely denote the 2 images. B) Electrodes are placed over the visual cortex; labeled according to the 10-20 International System (Cobb et al., 1958). C) We use CCA-filter on 8 electrodes to obtain a single spatially filtered time series EEG with good signal-to-noise ratios (Johnson and Krusienski, 2018).

EEG data were collected from 15 participants in total, each viewing two random sequences denoted as m-sequence 1 (**m-seq1**) and m-sequence 2 (**m-seq2**) for 2 trials each, given a total of 60 trials with a recording duration of at least 27 s for each trial (see **Methods** section for details). Four fixed-frequency trials were also performed for establishing characteristic SSVEP responses, which are not included in the present analysis.

In hopes of understanding the underlying physiology of the human visual pathway that manifests in EEG signals, we fit a transparent model to the EEG data. We decided to use relatively simple statistical time series models rather than a mechanistic differential equations model that would require many spatial scales (Young et al., 2019; Glomb et al., 2022) and specifying numerous (unknown) biophysical parameters. Aside from the absence of a good prescriptive model of the underlying system, another reason to use auto-regressive models is that Burg showed it is the maximum entropy model, i.e., makes minimally presumptive assumptions for capturing the autocorrelation of a single time series with lags up to the order of the model (Choi and Cover, 1984; Martini et al., 2024; Krusienski et al., 2006). We use the Box-Jenkins Methodology to fit these time series models to EEG, and to predict out-of-sample responses and validate our results (Shumway et al., 2000) (see **Methods** for more details).

### Stationarity Tests

The first step in the Box-Jenkins methodology is to perform a series of stationary tests of our EEG data to understand the benefits and drawbacks of the time series model fits. The tests used in this method are: Augmented Dickey-Fuller (**ADF**, tests against the null hypothesis that a given time series has a ‘unit root’, assuming the first coefficient *ϕ*_1_ has a *t* −distribution), Kwiatkowski-Phillips-Schmidt-Shin (**KPSS**, tests against the null hypothesis that a given time series is ‘trend-stationary’), and Phillips-Perron (**PP**, non-parametric version to ADF). In addition, Breusch-Pagan (**BP**) was used to test for homoscedasticity, i.e., the null hypothesis is homoscedasticity where the variance of the error/noise does not vary or depend on the variables (Breusch and Pagan, 1979; Koenker, 1981; Magris, 2024).

For all 60 trials, the ADF test and PP test resulted in the smallest *p*− value possible (***<*** 0.001), providing strong evidence that the EEG is stationary. The KPSS test resulted in the largest *p* − value possible (0.1) for all 60 time series, so we cannot rule out the null hypothesis that the data is stationary around a deterministic trend. In summary, all three of these tests provide some evidence that the EEG data are stationary.

The BP tests showed that some but not all of the EEG data were homeoscedastic, specifically with significance level *α* = 0.01, 32 out of 60 time series have *p*−values larger than significance so that the null hypothesis cannot be ruled out (28 are heteroscedastic); with *α* = 0.05, 40 out of 60 are heteroscedastic; see Fig A8 for further details.

### Analysis of EEG without Models

Before model fitting, estimation, and validation, we first provide a basic analysis of the EEG data without fitting a model. The power spectra for both m-seq1 (Fig 2A, top) and m-seq2 (Fig 2A, bottom) for all 30 trials (2 per each participant) in gray show some variability across participants and trials (the thick black curve is the population averaged power spectra across all 30 trials). In each of the 60 trials, the power spectra have sharp peaks at 3.5 Hz, with smaller values for larger frequencies, indicating that the EEG responses have a broadband but structured distribution for select frequencies. An additional peak, although much lower in magnitude, occurred at 18 Hz in both m-sequences. Since m-seq2 is m-seq1 in reverse order, the pseudo-random nature of the stimulus should prevent EEG responses from favoring one stimulus over the other. The average power spectra for m-seq1 and m-seq2-evoked responses were nearly identical (bottom row), supporting this assertion.

**Figure 2.**
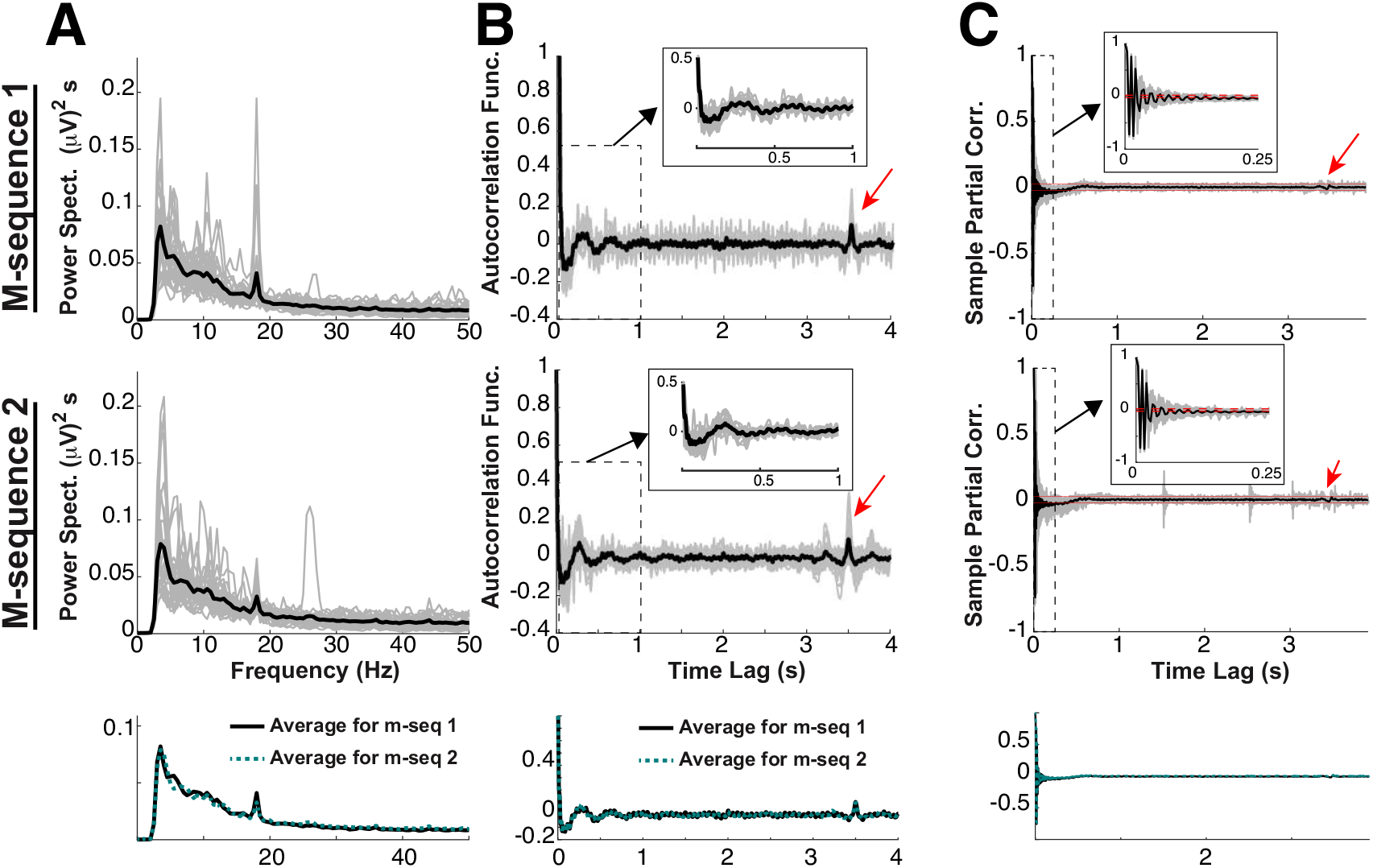
Summary of EEG response to two m-sequences (m-seq 1 in top row, m-seq 2 in middle row, bottom row compares the population averages). A) The power spectrum of EEG responses in (*µ*V)^2^*/*Hz. For all panels, the gray lines correspond to a single trial in one participant (30 total from 15 participants with 2 trials per stimulus lasting at least 27 s each); the black line is the average of all 30 gray lines. B) The autocorrelation function of the EEG responses; the dotted rectangle is a zoomed-in portion shown in the inset. The red arrow corresponds to the time-shift of m-seq input around 3.5 s to avoid steady-state stimulus frequency biases. C) The sample partial correlation function quickly decays and is relatively flat on average after moderate lag times. Here, the 30 red horizontal lines correspond to confidence intervals (1.96 std. dev.); note that they contain the population average (black curve) within less than 1 s. The middle panel of C (m-seq 2) shows gray curves that are visibly outside the error bars, corresponding to two outlier trials: participant 12 (trial 2), participant 6 (trial 2).

Autocorrelation functions (Fig. 2B) of the EEG responses oscillate at smaller time lags (***<***0.5 s) but tend to be suppressed for larger time lags, except for prominent peaks at around 3.5 s. Note that both m-seqs contained exact copies of the first 3.5 s of the sequence to avoid steady-state stimulus frequency biases (Waytowich and Krusienski, 2015), as evident with the peaks marked in Fig 2B,C with red arrows. Not surprisingly, all EEG responses contain statistically significant seasonal periods at 3.5 s because of the repeating input stimuli (with period 3.5 s). Partial autocorrelation functions (Fig. 2C) exhibit similar oscillations at small timescales, with consecutive time indices alternating between positive and negative correlation with the original EEG responses. The average partial autocorrelation across participants and trials reveals a slightly negative correlation, before stabilizing to 0 at around 0.7 s. Curves for larger time lags are within the confidence intervals (30 red lines barely distinguishable, 1.96 std. dev.) except for a few outliers. Seasonal behavior in the partial autocorrelation functions is also observed at 3.5 s, but at a smaller peak in correlation compared to Fig. 2B, indicating that smaller time indices are necessary to fully represent the seasonal aspect of the data. Both the autocorrelation and partial autocorrelation plots (Fig. 2B,C) do not precisely asympotote to 0 for longer time lags, indicating that a time series model consisting solely of an autoregressive without a moving average component may not capture all aspects of the EEG responses (Shumway et al., 2000). Similarly, a moving average model without autoregressive input would fail to capture the fast timescales of the EEG responses. Overall, this justifies considering time series models with both autoregressive and moving average components (ARMAX, SARMAX). Also, in some models, we include an exogenous input term to model the effects of the external stimuli (alternating frequencies) on the EEG responses. The long seasonal periods observed in Fig. 2B indicate that a seasonal component should also be included (see **Methods** for details).

### AR Suite of Models Fit to Data

Since there is some evidence that the EEG response data is stationary, we consider how well a suite of auto-regressive (AR) models fit the data, using a combined ordinary least squares (OLS), Yule-Walker (i.e., method of moments), and maximum likelihood estimation fit to 2 periods of data (in-sample, for 7 s, see Fig. A6A); we then test how well the fitted model predicts responses in the immediate period after (out-of-sample for 3.5 s). There were a total of 300 in-sample fits and out-of-sample estimates (150 for each m-seq; 5 intervals in each of the 4 total trials for each of the 15 participants). We chose a longer out-of-sample period than others (Pankka et al., 2025) because this enables good resolution of the frequency estimation (power spectrum). Although there are important challenges in accurately estimating the frequency content of short time segments of EEG that occur for example in brief tasks, due to lack of data and potential biases, which have been addressed by Birch et al. (1988); Vaz et al. (1987); Jansen et al. (2007); Krusienski et al. (2006).

Starting with the AR model, we systematically include an exogenous term (**X**), a moving average term (**MA**), and a seasonal term (**SAR**). We consider 4 total models (see **Methods: Model Structure** for details):

- AR 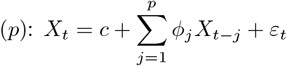, where *X*_*t*_ represents the EEG data, *ϕ*_*j*_ is the history kernel, ***ε***_*t*_ is uncorrelated Gaussian white noise, and ***c*** is the constant term.
- ARX 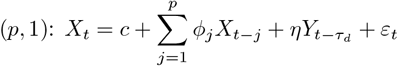, where *Y*_*t*_ is the exogenous external input obtained from the input.
- ARMAX 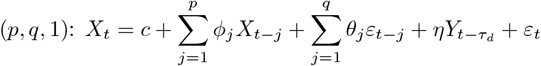, where *θ*_*j*_ are the moving average coefficients.
- SARMAX(*p, q*, 1, *P, s*): ARMAX with one (*P* = 1) multiplicative seasonal auto-regressive component of period length *s* = 896, or 3.5 s.

All models (except AR) had exactly one exogenous input term with a time delay of 70 ms (***τ***_*d*_ = 18) to model the signal traversing the visual pathway (Groen et al., 2022); alternatives using multiple lagged copies of the exogenous input did not fit the data as well (see **Methods** section ‘Model Order and Parameter Estimation’, second paragraph). In a given model, we determine the model order (*p, q*) by finding a candidate signal closest to the grand average of all 300 intervals and calculating the parameter (combinations) that gave the smallest AIC with maximum likelihood estimation, see Table 1.

**Table 1:**
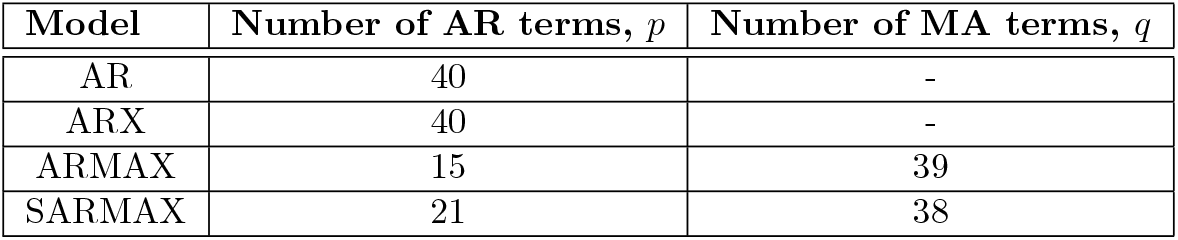
Fixed model parameter orders determined by the smallest AIC values.

| Model | Number of AR terms, $p$ | Number of MA terms, $q$ |
| --- | --- | --- |
| AR | 40 | - |
| ARX | 40 | - |
| ARMAX | 15 | 39 |
| SARMAX | 21 | 38 |

To implement the time-series model fitting to the EEG data, we have augmented several MATLAB functions (`estimate, simulate`) for several reasons: i) reduce the computational overhead and use only what we need, ii) handle rare cases when the model fits result in polynomial instability or non-convergence, in which case we use the next viable model fit, iii) to accurately start the out-of-sample estimation by using in-sample fits, accounting for when error terms ***ε***_*t*_ have history dependence. Further details are described in the **Methods: Parameter Estimation** section, and all of our new functions/scripts are freely available (see **Data Availability Statement**).

The in-sample fits of all 4 models are generally good despite some variability in the 300 total segments. For demonstration purposes, we show the ARMAX model fits, the results of which are displayed in Figure 3. The in-sample fits to all time-series segments for both m-seq1 and m-seq2 are good (lower right panel in Fig. 3A); the rest of Fig 3A shows the very best (top left) and very worst fit (bottom left) measured by the squared error averaged over the 1792 time points (7 s) which we refer to as the **mean-squared error**. Note that all (ARMAX) models only fit against statistically significant lag indices, informed by corresponding individual autocorrelation and partial autocorrelation functions in Fig. 2B and C. For example, all ARMAX models had at most 15 AR terms and 39 MA non-zero terms (Table 1), but their corresponding lag indices likely differed between participants and trials. This was done to use distant time indices for fitting while minimizing AR and MA polynomial complexity. The same approach was used for the other 3 models (Figs. A1–A3).

**Figure 3.**
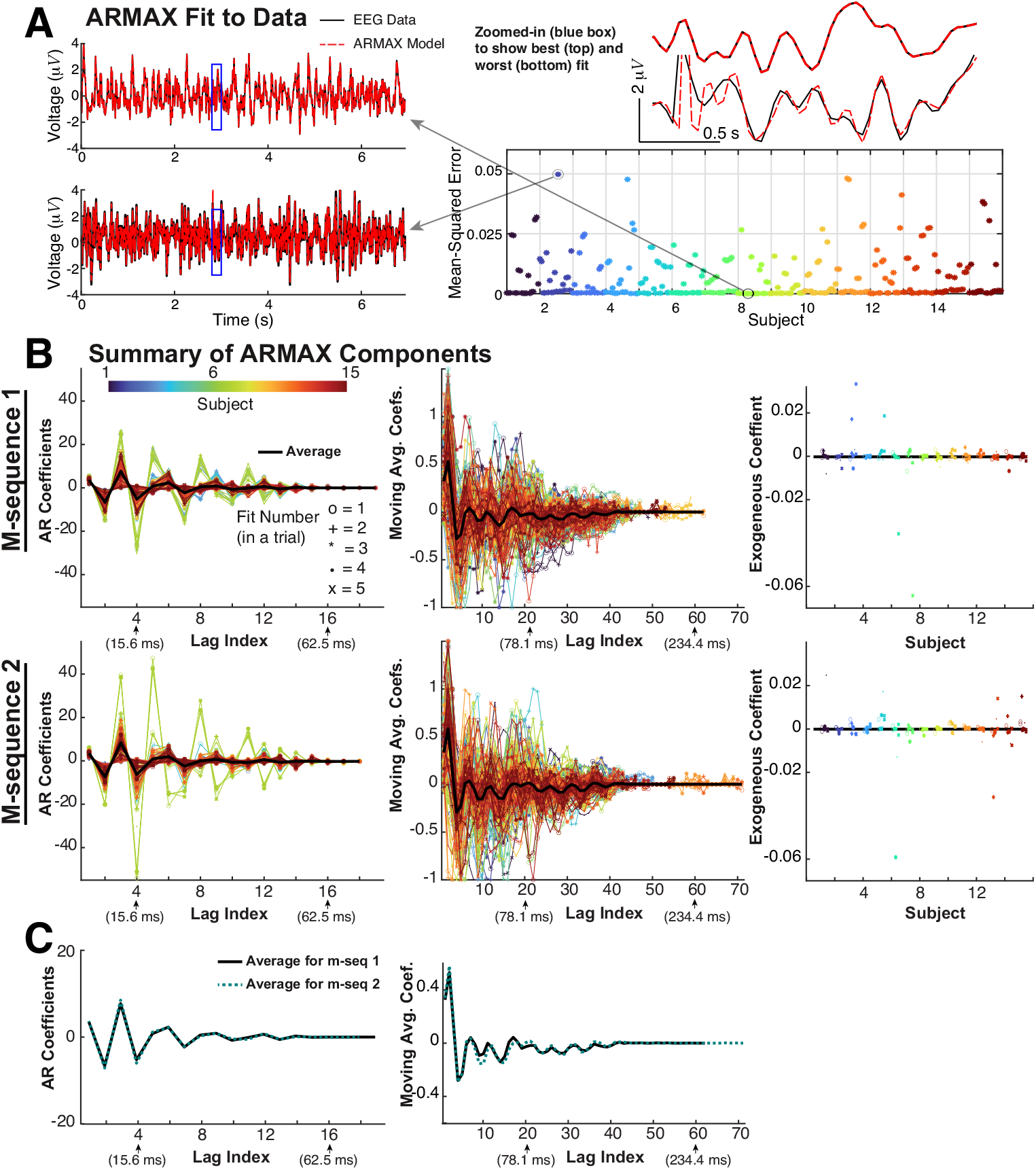
ARMAX fit to data. A) All the ARMAX model fits to EEG data are good overall. In a given interval we fit to 2 periods of EEG data (7 s) using a combined OLS, Yule-Walker, and MLE approach that amounts to a least-squares fit with a fixed number of AR coefficient terms, 1 term for the exogenous input, and moving average terms for all participants, trials, and intervals (see Eqn (3) and Methods for how we calculate the number of terms, and Table 1). In a trial, there are 5 different fits paired with out-of-sample testing for a total of 300 fits (15 participants, 2 trials each for both m-seqs) and thus 300 points in the lower right panel. B) ARMAX coefficient values from in-sample fits. C) Grand averages between both m-seqs are similar; the constant exogenous term has an average of −2.44 × 10^−4^ and −9.37 × 10^−5^ for m-seq1 and m-seq2 (resp.). Only values significantly different than 0 are plotted; some of the AR and MA terms are not shown at the end.

The in-sample fit results for all models clearly show some variability of model components, namely the AR coefficients, but variability is even more prominent in the MA coefficients for both ARMAX and SARMAX models. The AR coefficients in all models tend to oscillate between positive and negative values, with greater absolute values for a few participants (6 and 7, green curves). Another notable result is that for all 4 models, the grand average of AR and MA coefficients (when applicable) is quantitatively similar for both m-sequences (panel C). The exogenous terms (when applicable) exhibit some variability, but the values are overall relatively small (except for a few outliers in the SARMAX model, Fig. A3).

A truly impressive out-of-sample estimate would comprise a model capable of accurately fitting to point-wise EEG values. Currently, it is difficult for any EEG model to predict precise point-wise values, especially for 3.5 s on a fine temporal grid, so we instead aim to capture some statistical properties of the out-of-sample data that are known to be important (described in the next paragraph). We assess out-of-sample estimation performance of all 4 models by considering how well it captures EEG in the immediate 3.5 s period following the 7 s in-sample fits (Fig 4A). We do this for all possible participants, both m-sequences, all trials, and time intervals. Note that in each out-of-sample period, we simulate 1000 individual realizations of the model to adequately capture the average model behavior because there is a wide range of variability in a single noisy estimate. The following paragraphs will describe the metrics we use to quantify prediction performance (Fig 4), with a summary of the results shown in Figure 5 and Table 2; the granular details for all models and all 300 prediction periods are shown in the Appendix in Figs. A4–A7.

**Table 2:** Statistical test (Wilcoxon Rank Sum Test **WRST**) to assess whether out-of-sample estimation errors are different between models (see Fig. 5). Although there are some small *p*−values suggesting that we can rule out the null hypothesis that the errors/correlation are not from the same distribution, notice that all effect sizes are small with two exceptions; the effect sizes are *medium* (0.2, 0.5) or *large* (0.5, 0.8), a common qualitative comparison from Cohen (2013); Tomczak and Tomczak (2014).

| Relationship | Pt-wise Error |  | Pearson's Correlation |  | Power Spectrum Error |  |
| --- | --- | --- | --- | --- | --- | --- |
| | $p$-value | Effect Size | $p$-value | Effect Size | $p$-value | Effect Size |
| AR≠ARX | 0.97 | $1.4 \times 10^{-3}$ | 0.33 | $4.0 \times 10^{-2}$ | 0.98 | $1.2 \times 10^{-3}$ |
| AR≠ARMAX | 0.64 | $1.9 \times 10^{-2}$ | 0.99 | $2.4 \times 10^{-4}$ | $2.6 \times 10^{-4}$ | 0.15 |
| ARX≠ARMAX | 0.62 | $2.04 \times 10^{-2}$ | 0.30 | $4.2 \times 10^{-2}$ | $2.5 \times 10^{-4}$ | 0.15 |
| AR≠SARMAX | 0.23 | $4.9 \times 10^{-2}$ | $4.8 \times 10^{-2}$ | $8.1 \times 10^{-2}$ | $2.7 \times 10^{-7}$ | 0.21 (med.) |
| ARX≠SARMAX | 0.23 | $4.9 \times 10^{-2}$ | $3.0 \times 10^{-3}$ | 0.12 | $2.7 \times 10^{-7}$ | 0.21 (med.) |
| ARMAX≠SARMAX | 0.44 | $3.2 \times 10^{-2}$ | $4.3 \times 10^{-2}$ | $8.3 \times 10^{-2}$ | 0.056 | $7.8 \times 10^{-2}$ |

**Figure 4.**
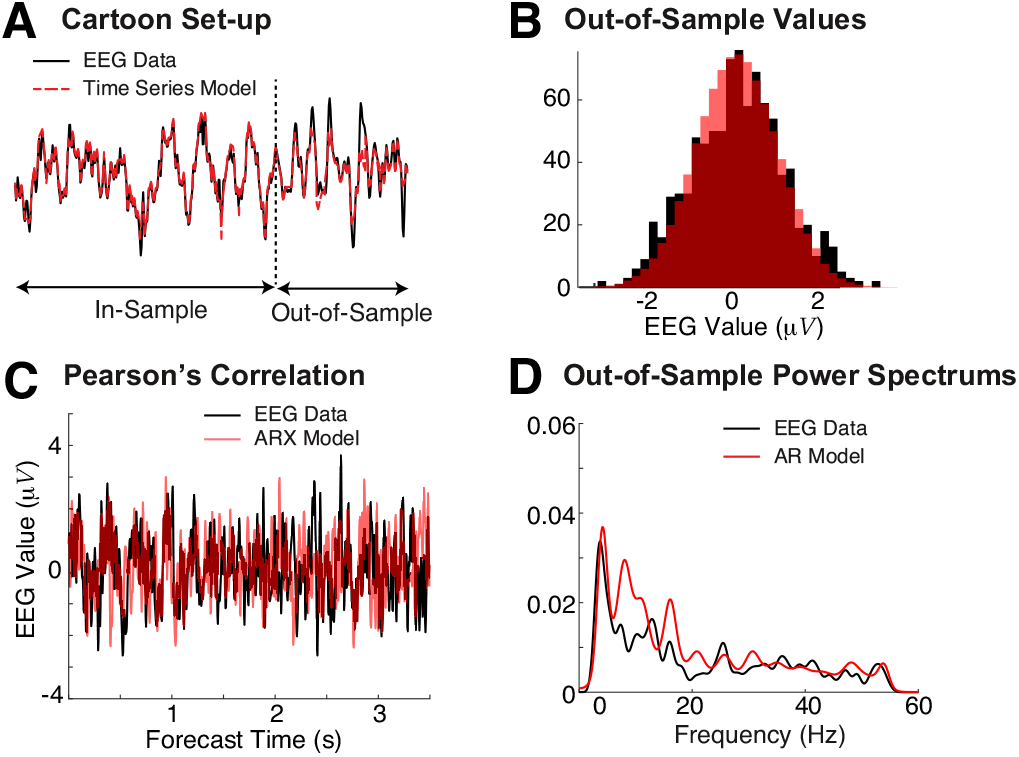
Quantifying quality of out-of-sample estimation. **A**) Cartoon of the set-up: the results of the in-sample fit (7 s) are compared to the immediate period (3.5 s) out-of-sample. **B**) In addition to calculating the mean-squared error (difference) between model and data, we also quantify the differences between the histograms of both model and data values (via mean-squared error). This example histogram is for the ARMAX model, corresponding to the average (out of all 300 out-of-sample estimates) mean-squared error of 29.3 (see Fig. A6B). **C**) Time series from an ARX model estimate and corresponding data: the Pearson’s correlation quantifies how the model and data move together in time. This example corresponds to the best Pearson’s correlation: 0.305 (see Fig A5C). **D**) Power spectra of both data and model in the out-of-sample period are calculated and compared using the mean-squared error. This is a good example of an out-of-sample estimate from the AR model with mean-squared error 2.2 × 10^−4^(see Fig A4Dii, interval 2 in dark blue).

**Figure 5.**
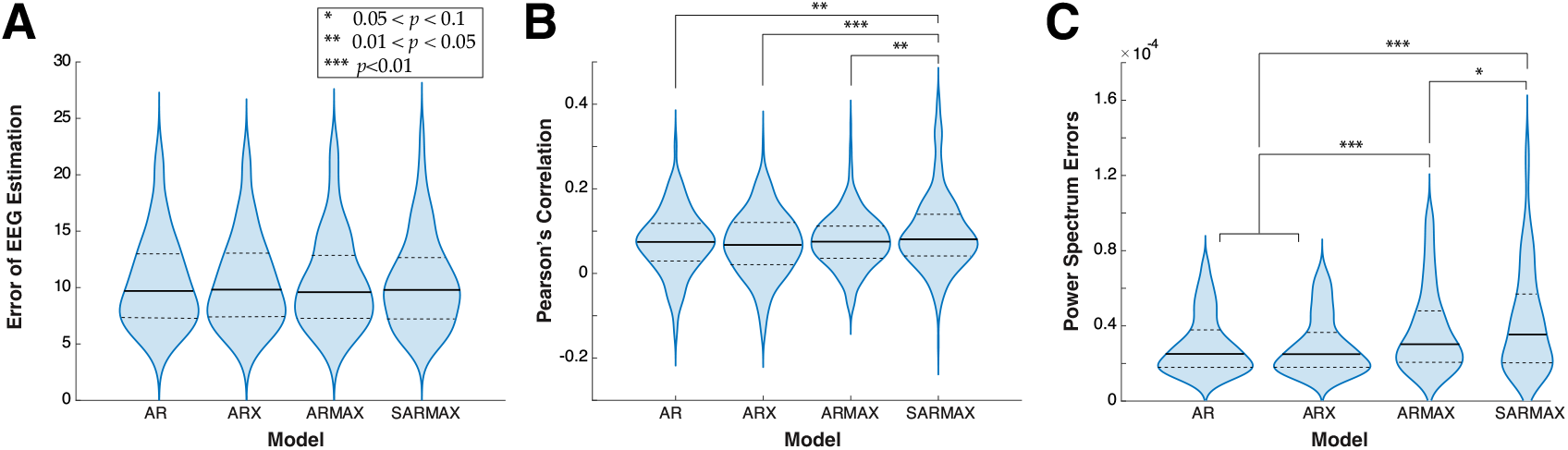
Violin plots comparing out-of-sample estimation performance for all 4 models. **A**) The distribution of the set of estimation errors (each point in a distribution is the average mean-squared error over the 3.5 s) is similar for all 4 models, with no statistically significant differences between any pair. **B**) The set of 300 Pearson’s correlations between data and model estimates is similar, the only difference is between the SARMAX model and the other 3 but with very small effect sizes. **C**) The distribution of mean-squared error between the power spectrum of the data and the model estimates can be statistically different: AR and ARX distributions of errors are similar, but they both differ from ARMAX and SARMAX; ARMAX and SARMAX are slightly different but all with very small effect sizes. Wilcoxon rank sum test was used to test differences, see Table 2. These violin plots excluded outliers for illustration purposes, but the statistical tests corresponding to the stars and Table 2 were performed on the full dataset for all models.

We first consider: i) histogram of values in the out-of-sample estimation period (3.5 s period out-of-sample immediately following the in-sample period) and analyzing the mean-squared error (difference) between data and model (Fig. 4B; see Figs A4–A7B for individual models), ii) the Pearson’s correlation in time between the actual EEG and realization-averaged out-of-sample estimates (Fig 4C; see Figs A4–A7C for individual models). The histograms of the point-wise time series are a coarse measure absent of the temporal dynamics (but see Fig 5A), but the suite of AR models can perform well by this measure. In fact, the average (mean) of all mean-squared errors is small. Pearson correlation values were measured against the average out-of-sample estimates as opposed to their individual realizations since averaging removes instances of white noise and does not capture the range of Pearson’s correlation values across realizations, giving a linear correlation between a representative prediction and the EEG data. As a result, the Pearson correlation values are generally positive (see right most panel in Figs A4–A7C) but the performance is variable, as evident with time series plots for the entire out-of-sample period for the best, mean, and worst overall correlation.

The frequency content of EEG responses is a crucial entity in many scientific and clinical studies (Newson and Thiagarajan, 2019), including associations with different brain states, sensory perception, cognitive performance, etc. (Buzsáki, 2006), so we consider how well the models can capture individual EEG power spectra in various out-of-sample periods (Fig 4D; see Figs A4–A7D for individual models). Details in the Appendix show for each model and each m-seq comparisons of data and model for all 5 prediction intervals within a trial that had the best and worst average (over 5 segments) mean-squared error; we also show in the right-most panel the mean-squared errors for all prediction intervals (Figs A4–A7D).

We see in Figure 5 and Table 2 that overall, there is surprisingly little difference in out-of-sample estimation performance between the 4 models. We use the Wilcoxon Rank Sum Test (**WRST**) to compare various pairs, with the null hypothesis that the 2 sets are drawn from the same distribution: we report *p*−values and effect sizes (see **Methods**) in Table 2. Despite some statistically significant differences in the distribution of errors (or Pearson’s correlation values), the effect sizes were generally small (Table 2) with two exceptions: power spectrum errors between AR/ARX and SARMAX had medium effect sizes. The power spectra of the out-sample estimates in all 4 models had some variation, with AR and ARX performing best, followed closely by ARMAX and then SARMAX (Fig. 5C). We conclude that, for the most part, quantitative differences in estimation performance measures are minor except when considering frequency content (power spectra).

We also consider how well the models predict EEG in different frequency bands, specifically using 4 common frequency bands: *δ*: [1,4] Hz, *θ* : [4,8] Hz, *α* : [8,13] Hz, *β* : [13,30] Hz (Buzsáki, 2006; Newson and Thiagarajan, 2019). We assess this in 2 ways: i) within a given frequency band, we determine whether the 4 models are different, and which, if any, perform better; ii) within a given model, we determine whether the errors are different between frequency bands. Here, we simply examine a sub-domain of frequencies in the desired band, calculating 300 mean-squared error values between the power spectrum of the out-of-sample estimates and the actual power spectrum from EEG data.

Comparing all models within a given frequency band (Fig. 6, Table 3), we see that AR and ARX have smaller errors than ARMAX and SARMAX in delta- and theta-bands, and they (AR and ARX) have smaller errors than SARMAX in the beta-band. There is surprisingly no difference between any pairs in all 4 models in the alpha-band. Within all frequency bands, ARMAX is different than SARMAX.

**Table 3:** Within a frequency band, assessing differences in out-of-sample estimates of the power spectra across models using WRST, reporting the *p*−values and effect sizes.

| | Relationship | AR≠ARX | AR≠ARMAX | AR≠SARMAX | ARX≠ARMAX | ARX≠SARMAX | ARMAX≠SARMAX |
| --- | --- | --- | --- | --- | --- | --- | --- |
| $\delta$ | $p$-value | 0.98 | 0.07 | $2.2 \times 10^{-4}$ | 0.07 | $2.2 \times 10^{-4}$ | 0.06 |
|  | Effect Size | 0.001 | 0.07 | 0.15 (small) | 0.07 | 0.15 (small) | 0.08 |
| $\theta$ | $p$-value | 0.99 | 0.002 | $2.9 \times 10^{-6}$ | 0.002 | $3.3 \times 10^{-6}$ | 0.08 |
| | Effect Size | $5.7 \times 10^{-4}$ | 0.13 (small) | 0.19 (small) | 0.13 (small) | 0.19 (small) | 0.07 |
| $\alpha$ | $p$-value | 0.96 | 0.99 | 0.58 | 0.95 | 0.56 | 0.59 |
| | Effect Size | 0.002 | $5.1 \times 10^{-4}$ | 0.02 | 0.003 | 0.024 | 0.02 |
| $\beta$ | $p$-value | 0.93 | 0.82 | 0.009 | 0.76 | 0.007 | 0.02 |
|  | Effect Size | 0.004 | 0.009 | 0.11 (small) | 0.01 | 0.11 (small) | 0.10 (small) |

**Figure 6.**
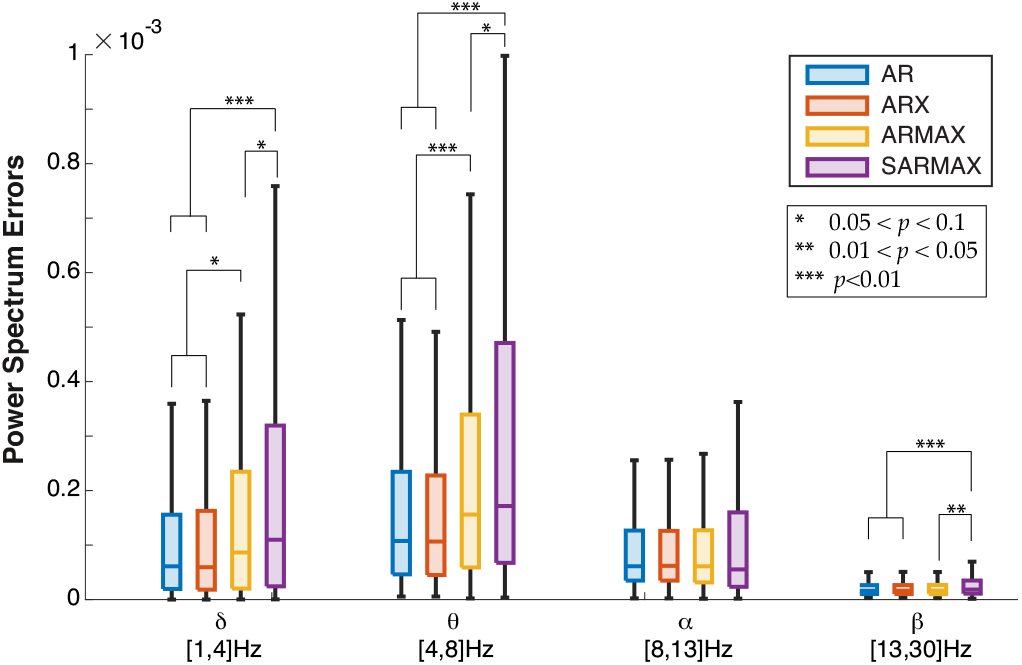
Comparison of out-of-sample power spectrum errors by frequency bands. Using WRST, we find both AR and ARX models have different distributions of mean-squared errors than SARMAX in the delta-, theta- and beta-band (all bands except alpha) with varying levels of statistical significance but generally small effect sizes (see Table 3). In the theta-band, AR and ARX are also different than ARMAX. In the beta-band, ARMAX and SARMAX are slightly different. In the alpha-band, there are no statistically significant differences between models.

For a given model, the average performance is best in the beta-band, followed by alpha-, delta-, then theta-, but whether these differences are statistically significant is unclear. Again we use WRST (Table 4) to find that the difference between frequency bands are generally statistically significant with reasonable effect sizes (mostly medium to large, a few small); the only exceptions are that the delta- and alpha-bands are not different in all models except SARMAX (with small effect size). Overall, the power spectrum of the out-of-sample estimates does depend on the frequency band of interest.

**Table 4:** Differences in out-of-sample estimation errors across frequency bands within a given model using WRST, reporting the *p*−values and effect sizes.

| | Relationship | $\delta \neq \theta$ | $\delta \neq \alpha$ | $\delta \neq \beta$ | $\theta \neq \alpha$ | $\theta \neq \beta$ | $\alpha \neq \beta$ |
| --- | --- | --- | --- | --- | --- | --- | --- |
| AR | $p$-value | $1.0 \times 10^{-7}$ | 0.50 | 0 | $8.2 \times 10^{-8}$ | 0 | 0 |
|  | Effect Size | 0.22 (med) | 0.03 | 0.44 (med) | 0.22 (med) | 0.71 (lrg) | 0.58 (lrg) |
| ARX | $p$-value | $8.7 \times 10^{-8}$ | 0.42 | 0 | $8.3 \times 10^{-8}$ | 0 | 0 |
|  | Effect Size | 0.22 (med) | 0.03 | 0.44 (med) | 0.22 (med) | 0.7 (lrg) | 0.58 (lrg) |
| ARMAX | $p$-value | $1.2 \times 10^{-7}$ | 0.19 | 0 | $2.0 \times 10^{-15}$ | 0 | 0 |
|  | Effect Size | 0.22 (med) | 0.03 | 0.44 (med) | 0.22 (med) | 0.70 (lrg) | 0.58 (lrg) |
| SARMAX | $p$-value | $8.4 \times 10^{-6}$ | 0.004 | 0 | $8.4 \times 10^{-16}$ | 0 | 0 |
|  | Effect Size | 0.18 (small) | 0.11 (small) | 0.41 (med) | 0.33 (med) | 0.62 (lrg) | 0.38 (med) |

## 3 Discussion

We collected EEG data with electrodes placed over the visual cortex while participants viewed spatially localized stimuli, alternating at pseudo-random temporal frequencies. We systematically fit several statistical time-series models using the Box-Jenkins methodology, where maximum likelihood fits and AIC determined the number of parameters. Optimal least squares and Yule-Walker (method of moments) were subsequently used to fit models to data. We fit each model to two periods of random input (7 s) and tested the prediction power of the model in the immediate out-of-sample period for 3.5 s, assessing the differences between model and data via the point-wise values, histograms of values, Pearson’s correlation between model and data, and differences in the power spectrum of data and model. We find that the set of auto-regressive models with varying complexity, i.e., possibly with exogenous input, moving average terms, seasonality component, generally did not vary much in out-of-sample estimation performance. The power spectra of the out-of-sample model estimates were different depending on the model, and there were differences in frequency bands; simple auto-regressive models performed better than the other models on average, and the beta-band (followed by alpha-, delta-, then theta-) was where the out-of-sample power spectrum was most accurate for all models. The time series models we use have a long history of usage for many complex time series (Shumway et al., 2000), including EEG (Vaz et al., 1987; Birch et al., 1988), but here we have collected new data and compared various types of time-series models. This study presents a viable framework to fit other EEG data using time-series models, providing insights to the human EEG response that could generalize to other visual stimuli.

We use common stimuli in brain-computer interface (BCI) studies, m-sequence stimuli with pseudo-random temporal frequencies, to elicit complex temporal EEG responses. Stimuli alternating with fixed temporal frequencies are commonly (**?**) and still currently used in BCI (**?**), whereas m-sequences have much richer temporal dynamics and certain advantages in the context of BCI (Bin et al., 2009). It is unclear whether such simple time-series models are effective beyond such structured temporal stimuli. More complex visual stimuli, such as arbitrary images or natural scenes, would require the incorporation of spatial information into the models. One ambitious direction for future research is to develop a more principled detailed biophysical model that produces reliable and accurate EEG for a broader range of visual input stimuli.

This study does not make any general claims about the physiology of the human visual system, rather we only conclude how to capture some aspects of the EEG response with the specific m-sequence inputs considered. Since the auto-regressive terms oscillate between positive and negative values for all models and subjects (Fig 3, A1–A3), taken together with the relatively good performance of the simple AR model suggests that the weighted history of EEG values is crucial for estimating this data with these models. One interpretation of these results is that any such model should have history dependence spanning at least 50 ms up to 100’s of milliseconds, i.e., where the auto-regressive terms decay to near 0. That is, the effective timescales of a mechanistic model of EEG should be in this range and should have oscillations so that the model’s values depends on past history. Whether this holds in general for other stimuli is unclear, but such an investigation would require similarly robust results with perhaps a more general model.

Curiously, the SARMAX model does not improve out-of-sample performance compared to the other simpler time series models. In adding a seasonal component, a very long lag wherein model results from 3.5 s previously (896 terms before) contribute to the model at a given time point, but note that the in-sample fits yield a relatively small seasonal term (Fig A3) that would therefore not improve performance. The SARMAX model was our attempt to capture the modest periodic component observed in the ACF of the EEG at lag 3.5 s (Fig 2B). This does not rule out the possibility of the existence of another with a meaningful long seasonal component that results in a significant improvement to the other time series models (AR, ARX, ARMAX).

A dichotomy of approaches to understanding new neural data broadly consists of mechanistic models that aim to uncover underlying neural attributes that drive experimental observations, or statistical models that are principally fit to data to have the most accurate out-of-sample estimates at the expense of abstracting away core neural attributes (Perretti et al., 2013). We opted for the latter approach because EEG’s complex dependence on many smaller-scale neural circuit components (Pesaran et al., 2018) (and propagation of electrical signals through gray matter, skull, and scalp) makes developing viable and useful mechanistic models of this whole pathway difficult. A realistic biophysical model of this extended visual pathway from visual stimuli to EEG is challenging despite numerous animal studies that have led to advances in our knowledge about the underlying physiology. EEG data is noisy with a relatively low SNR compared to direct spiking or other anatomically invasive neural voltage recordings. We would be remiss without mentioning some of the ambitious works that address this issue, for example Neymotin et al. (2020); Tolley et al. (2024) connect EEG to multi-compartment pyramidal neural networks when there is a a strong input; the popular dynamic causal modeling (Kiebel et al., 2008) infers differential equations to describe interaction of different brain regions that best explain spatial data. The best mechanistic models fit to data generate a set of predictions about the underlying neural attributes that may then be validated with non-human mammalian experiments, but all of this is beyond the scope of this current study; future studies warrant endeavoring on this approach.

There are numerous non-mechanistic black-box models that rely extensively on fitting to ample data to obtain biological insights, with many contemporary studies using deep neural networks (Pathak et al., 2022; Li et al., 2020) and some paired with sophisticated Bayesian frameworks where predictions can account for uncertainty in the prediction (Pankka et al., 2025). A natural starting point for our new data are AR models and their variants (ARX, ARMAX, SARMAX) because they are simpler with model components that are easier to understand. The components of the model have a clear role in shaping the outputs, and fitting these time-series models results, in all cases, convergence to at worst a local minimum (at best a global minimum) and a causal, polynomial stable model, at least in this dataset and within the framework here. Also, we use relatively fewer parameters and degrees of freedom compared to contemporary deep neural network models; although with complicated visual scenes that are more spatially rich than our stimuli might require more complex models with more parameters.

A recent study (Pankka et al., 2025) compared how well AR models perform compared to a deep neural network model WaveNet, although they used a larger EEG dateset (68 subjects, 60 channels) collected from humans learning to perform a task with reward and punishment (Cavanagh et al., 2019), and WaveNet was developed to generate waves corresponding to audio (Van Den Oord et al., 2016). Nevertheless, Pankka et al. (2025) found that WaveNet outperformed a basic AR model in predicting signal amplitude and phase of EEG in the theta and alpha band. As already mentioned, there are many material differences between that study and ours (i.e., our focus on random temporal frequency input, time windows for fitting and predicting out-of-sample responses are different, etc.); a fair comparison between WaveNet or other deep neural network models with the time-series models here would be an interesting future study, but is beyond the scope of this current paper.

## 4 Methods

### EEG Data

Informed written consent was obtained from all participants for being included in the study. This study was approved by the Institutional Review Board of Virginia Commonwealth University according to the principles expressed in the Declaration of Helsinki.

EEG data was collected from 15 healthy participants with either 20/20 vision or corrected to normal vision with glasses/contacts/lasik, consisting of 8 males and 7 females, with ages ranging between 22–59 years old (mean age: 31, std. dev.: 12). Participants who usually wear corrective eye wear (glasses, 7 total) did not do so during experiments because of VR headset discomfort, but distinction between light and dark regions and spatial fidelity were still visible to all participants due to the simplistic nature of the stimuli. We first presented this data in Delaney (2023). The study was reviewed and approved by the Institutional Review Board of Virginia Commonwealth University.

Participants were instructed to focus on the visual stimulus consisting of alternating patterns of 2 concentric circles with white/black checkered patterns in the center of the receptive field where white → black and black → white (Fig. 1A). The visual stimulus was set to appear 60 cm away in virtual space and scaled to have a total diameter of 10 cm, covering 7.3° visual angle to all participants. This spatial frequency was set to elicit strong EEG signals (Waytowich et al., 2016) (Chapter 5.2 in Delaney (2023)). To ensure participants maintained focus, in each trial, participants were instructed to press a button when a red dot randomly appeared at the center of the stimuli; the EEG data during this attention test was removed from our analysis. Participants also took self-paced breaks between sessions as needed.

In total each participant viewed 6 different flickering stimuli: four with fixed frequencies of: 5.625, 6.429, 7.5, 9 Hz, and 2 pseudo-random m-sequences lasting about 27 s for each trial, sampled at 256 Hz with time bin length 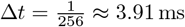. There were 2 trials of each of the 6 different stimuli for a total of 12 sessions for each of the 15 participants, but our modeling focused entirely on the m-sequences. The fixed frequencies are similar to those used in SSVEP-BCIs (steady state visually evoked potentials in brain-computer interfaces), and the m-sequences are an established protocol used in cVEP-BCIs (coded visually evoked potentials in brain-computer interfaces).

Eight electrodes were positioned over the occipital region, with locations based on the International 10-20 system (Cobb et al. (1958), Fig. 1B). In the fixed frequency experiments, a set of 3 sine and 3 cosine functions, each with 1 driving frequency and 2 harmonic frequencies of the visual stimulus, constituted a template set of signals for which canonical correlation analysis (CCA) was applied to determine the weights (spatial filtering) that gave the largest correlation with the (linearly transformed) template set (Fig 1C).

For the 2 m-sequences, a similar procedure was used but the average EEG response to the visual stimulus constituted the entire template set since they have a large frequency bandwidth. This heuristic is widely adopted and established to be effective for classification results (Johnson and Krusienski, 2018; Bin et al., 2011). Like in the fixed frequency case, the linear combination of the canonical weights and the individual electrode responses form a singular, spatially-filtered EEG signal, but it is maximally correlated with the average EEG response across electrodes instead of a sinusoidal template set. See (Delaney, 2023) for additional information on electrode placements and CCA of EEG responses during m-seq trials.

### Box-Jenkins Methodology

Below we provide a brief checklist of the Box-Jenkins methodology, which is a standard approach to constructing autoregressive models for time series data (Shumway et al., 2000).

- **Stationary Testing:** Time series data must be stationary before attempting to fit the data with a model. Seasonal and non-seasonal differencing or detrending are effective tools to force stationary means, but ADF, KPSS, PP test results indicated no data transformations were required in the present data set.
- **Model Order Selection:** Autocorrelation (ACF) and partial autocorrelation function (PACF) plots are typically employed to determine moving average and autoregressive component orders, respectively. If index significance cuts off at a specific lag, then the component order can be fixed at the cut-off, but when both ACF and PACF plots decay over long periods, as in the present case, then an ARMA model is deemed minimally necessary in theory. Out-of-sample estimation performance ultimately determines model order and structure, which is left to the modeler. An alternative approach involves the use of metrics such as the Akaike (AIC) to identify the optimal model order by exhaustive search.
- **Parameter Estimation:** Ordinary least squares (OLS), Yule-Walker (method of moments), and maximum likelihood estimation (MLE) techniques are popular methods of estimating model parameters, which include but are not limited to the autoregressive, moving average, and exogenous coefficients and the constant and innovation distribution variance values. Estimation procedures are typically predetermined by the employed software package. We use a combined method to estimate model parameters, with more details listed below.
- **Model Validation/Prediction:** The Ljung-Box Q test or any comparative test determines if in-sample residuals are white noise or serially correlated. The presence of serial correlation suggests the model is not complex enough to capture certain temporal structure found in the data and warrants model reconstruction. If residuals are white noise correlated, then the model is fit for predicting out-of-sample. As a result, it generally informs on whether the augmentation or elimination of model components benefits the model’s ability to capture structures found in data, but since we are comparing various autoregressive-based models, out-of-sample estimation performance would be a better metric to measure model strength.
- **Prediction Performance:** It is common practice to compare out-of-sample prediction accuracy across different autoregressive model types to inform suitable models of inference. Mean absolute error (Pankka et al., 2025), mean squared error (MSE), Pearson’s correlation, and other metrics are commonly used to quantify this. Lower mean absolute and mean squared errors and higher Pearson’s correlation values indicate higher out-of-sample estimation accuracy and determine an optimal predictive model among possible candidate models.

## Model Structure

We use a suite of autoregressive models, starting with the simplest AR model and systematically augmenting with exogenous, moving-average, and seasonality components.

### AR Model

The initial AR model was used to model EEG data across all m-sequence trials. The AR(*p*) model of order *p* is defined as:

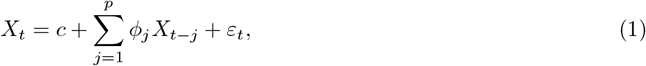

where *X*_*t*_ represents the EEG data, *ϕ*_*j*_ is the autoregressive coefficient at lag index *j, ε*_*t*_ is an instance of Gaussian white noise with variance *σ*^2^ at time *t*, and *c* is the model constant term. AR(*p*) models commonly use lag indices up to and including the *p*^th^ index to model a stationary process, but doing so risks higher model complexity in time series with longer timescales. Our AR model only considers statistically significant lag indices (*α* ***<*** 0.05), informed by the partial autocorrelation function. We define model order as the number of terms present in a given process, to avoid confusion related to discrepancies in autoregressive lag indices across participants and trials. The constant term ***c*** and variance ***σ***^2^ are simply the mean and variance of the EEG data, assuming the data were normal and independent of time. Details on estimating the history kernel {*ϕ*_*j*_} are presented in **Parameter Estimation**.

### ARX Model

The autoregressive model with exogenous input (**ARX**) is obtained from augmenting the AR model in Eq.1 with an exogenous variable, *Y*_*t*_. The subsequent ARX(*p*, 1) model of autoregressive order *p* with one exogenous input variable is defined by the following equation:

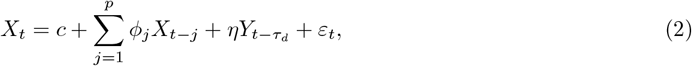

where 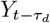 models the input stimulus. *Y* is obtained from the stimuli by mapping the images to a corresponding binary sequence: 0 for an image and 1 for reversed image (white → black and black → white), then low-pass filtering to model neural network integration of inputs along the visual pathway (Hawken et al., 1996) using the MATLAB 2024b command lowpass with a threshold of 9 Hz and sampling frequency of 256 frames per second. Moreover, the exogenous variable 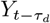 has an effective time delay of 70 ms, or approximately ***τ***_*d*_ = 18 lag indices, to account for the time for the signal to traverse from the retina to the primary visual cortex (Groen et al., 2022).

### ARMAX Model

The autoregressive, moving-average model with exogenous input (**ARMAX**) was constructed by augmenting the ARX model in Eq.2 with a moving average process of order *q*, resulting in the following ARMAX(*p, q*, 1) model:

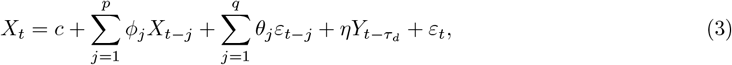

where *θ*_*j*_ represents the moving average coefficients and *ε*_*t*−*j*_ represents error terms or white noise instances, depending on whether the index *t* − *j* lies within in-sample or out-of-sample periods. The variance *σ*^2^ of the white noise process *ε*_*t*_ was still estimated as the variance of the data. Similar to AR and ARX models, the moving average process *MA*(*q*) only considers statistically significant lag indices (*α <* 0.05), informed by the autocorrelation function, as a means of reducing model complexity while regressing over long timescales. Due to our characterization of model order, an ARMAX(*p, q*, 1) model has *p* autoregressive terms *X*_*t*−*j*_, *q* moving average terms *ε*_*t*−*i*_, and one exogenous input term 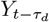. An equivalent alternative form for Eq. 3 is obtained through the use of backshift operators, where 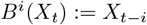 for any positive integer *i*:

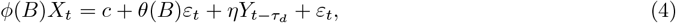

where 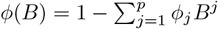 and 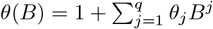 are the corresponding autoregressive and moving average backshift polynomials.

### SARMAX Model

Long seasonal periods in recorded EEG autocorrelation plots (Fig. 2B) suggested that we should consider candidate models with a seasonal autoregressive process, i.e., the seasonal autoregressive, moving-average model with exogenous input (**SARMAX**). This was the most complex model under consideration. Regressing over a seasonal period means including not only the function value at index *s* but also the previous time indices from index *s* up to *r* + *s*, where *r* is the maximum time lag used in the non-seasonal autoregressive process. Regressing over both seasonal and non-seasonal indices is best expressed in closed form by multiplicative seasonal and non-seasonal backshifting polynomials:

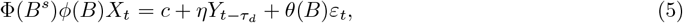

where 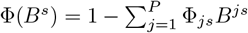 and 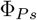 represents the seasonal autoregressive coefficient with period length ***s*** and seasonal index *P* . We allowed our SARMAX(*p, q*, 1, *P, s*) model to regress against one seasonal period (*P* = 1) of length *s* = 896, simplifying to the following considered model:

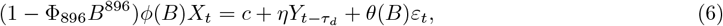

### Parameter Estimation

Note that all new scripts/functions we created to handle this dataset are freely available; see **Data Availability Statement**.

#### Order Selection

The Box-Jenkins methodology states that PACF and ACF plots can determine corresponding autoregressive and moving average component orders. If the PACF has sudden decreases in significance at lag *p*, then an AR model of order *p* is sufficient to model the time series. Likewise, for an MA model of order *q* when the ACF cuts off at lag index *q*. However, when both functions decay after respective lag indices *p* and *q* with a non-evident cut-off, an ARMA(*p,q*) model is appropriate. The specific selection of orders *p* and *q* in this scenario are often chosen arbitrarily within a region of possible values unless informed by another method. To ensure that model order selection is not exclusively left to the researcher, we used an exhaustive AIC search with model fit to a candidate signal to obtain an optimal model order across all participants and trials. The candidate signal was selected as the signal closest (in a Euclidean manner) to the average EEG signal across all participants, trials, and segments. AR and ARX models of the candidate signal (participant 9, segment 1, trial 1 over m-seq 2) were estimated with an autoregressive term count ranging from 1 to 90. ARMAX models were estimated on the candidate signal across an exhaustive grid of 0 to 60 autoregressive and moving average terms, and SARMAX models were estimated on a grid of 0 to 39 autoregressive and moving average terms. A more restrictive search was selected for SARMAX entirely due to the extremely high computational cost. Orders for each model type were selected by the parameter combination that resulted in the smallest AIC value. We mention again that while model orders within model type are conserved across all participants and trials, models only fit non-seasonal autoregressive terms and moving average terms using statistically significant lag indices from their respective PACF and ACF plots. Hence, while the number of autoregressive and moving average terms will be constant across participants and trials within a model type, the specific lag indices under regression are subject to change depending on the participant and trial. Exogenous variable order was fixed at one ***η*** term in ARX, ARMAX, and SARMAX models for two reasons: 1) a coarse-grained search for the lowest AIC value across various exogenous input orders resulted in an optimal value of 1 term and 2) models were constructed to quantify the impact of a single present stimulus on a spatially-filtered EEG signal. From a modeling standpoint, including any further time-dependent exogenous variables increases the risk of model instability and non-stationary out-of-sample estimates. Optimal component orders for all model types are in Table 1.

#### Data Partitioning

For all AR, ARX, and ARMAX models, EEG data were split into five forward-shifting overlapping intervals of 2688 data points (3 periods of 896 data points or 3.5 s), with each interval being time base partitioned into the following three consecutive periods: 1) a presample period, 2) an in-sample training period, and 3) an out-of-sample esimtation period. The presample period includes data samples up to the maximum lag index in the model to initialize lagged component values. As such, data in the presample period were not used directly in estimating model parameters. For all SARMAX models, EEG data were split into intervals of 3584 data points (4 periods of 896 data points) to accommodate an increase in presample period length of 896 data points while keeping in-sample and out-of-sample period lengths consistent across model types for estimation and model comparison purposes. Small differences in in-sample period length were observed across participants due to the significance of specific lags in corresponding ACF and PACF plots (see Fig. 2).

#### Estimation Algorithm

We estimated model parameters across 300 intervals of EEG data (15 participants, 4 trials per participant, 5 intervals per trial) by using a combined OLS, Yule-Walker, and MLE approach. The estimation procedure was numerically implemented by augmenting the MATLAB function estimate to remove post-estimation steps, creating new m-files: `estimate_mod/` and `estimate_mod_hist`, to speed up computations; this also uses `fmincon` to solve a constrained nonlinear optimization problem on the following log-likelihood function, assuming a Gaussian innovation distribution:

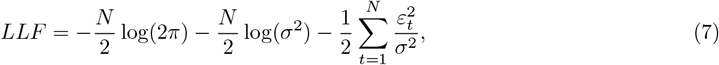

where *N* represents the in-sample period length, *σ*^2^ is the innovation variance, and *ε*_*t*_ is the innovation process. Initial estimates for all parameters were obtained using an OLS technique, except for the moving average coefficients in the ARMAX model, which were prescribed by solving the modified Yule-Walker equations. Convergence criteria and optimization options were selected to ensure that SARMAX model parameters could be obtained reliably, given that SARMAX was the most complex model under consideration. Accordingly, the optimization problem used a sequential quadratic programming-based (SQP) algorithm with a maximum function evaluation count of 1e10, maximum iteration count of 2e3, and step tolerance of 1e 10. In the rare instance of non-convergence, the last feasible parameter set in the algorithm was designated as optimal, making use of our new function `mod_hist`. When there was failure to converge using the SQP algorithm, we used a simulation repeat using the SQP-legacy algorithm (see new scripts `estimate_mod` and `estimate_mod_hist`). Autoregressive and moving average coefficients were constrained such that their corresponding backshift polynomials would only have roots lying outside the unit circle in the complex plane. This was done to ensure that model predictions were stable and not overly reliant on past errors. In-sample residuals between estimated models and EEG data were generated using the MATLAB function `infer`, which infers errors, i.e., instances of the innovation process *ε*_*t*_, at each point in time when the model is iteratively fitted to every individual data point in the in-sample period. In-sample model fits were then generated by adding the residuals back to the in-sample EEG data. The mean squared error between the in-sample model fit 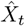 and EEG data *X*_*t*_ was collected to quantitatively measure the quality of fit.

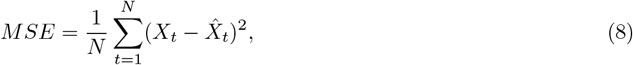

where *N* represents the in-sample period length.

### Out-of-sample Estimates

Simulated realizations of all estimated models were produced by augmenting the function `simulate` to use model values from the in-sample period (by default `simulate` does not do this); we created new functions `simulate_mod` and `simulate_mod_hist` to predict out-of-sample responses. We generated 1,000 individual model realizations when supplied with corresponding in-sample data and in-sample and out-of-sample exogenous variable data. For ARMAX and SARMAX models, in-sample error terms *ε*_*t*−*j*_ were supplied by the in-sample residuals, whereas error terms indexed in the out-of-sample period were generated by a Gaussian innovation distribution with zero mean and constant variance *σ*^2^, which was obtained during parameter estimation. For AR and ARX models, which lack a dependence on past errors, all error terms *ε*_*t*_ were simply generated by the innovation distribution.

Prediction performance was measured quantitatively according to the following two metrics: 1) the mean-squared error of EEG data and histograms of individual realizations, and 2) the Pearson’s correlation of EEG data and the realization-averaged prediction. Other measures of prediction accuracy, such as the mean-squared error in time of the average out-of-sample estimates and the EEG data, were not considered primarily because of the relatively fast decay of average estimates observed over a long interval shared across model types. Such a metric would fail to capture small-scale differences in the average estimates between model types. The Pearson’s correlation of the averaged estimates and EEG data would be a better metric since it quantifies the linear relationship instead of the fit quality.

Histograms partitioned individual realization values and EEG data in the out-of-sample period into *N*_*hist*_ = 100 bins with values ranging from -10 to 10 at edge intervals of 0.2. The mean squared error of the EEG data bin count and the bin count averaged across individual realizations was measured according to the following formula, given that *H*_*i*_ represents the bin count of the EEG data and 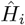 represents the average bin count across individual realizations:

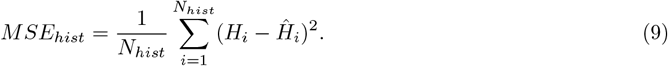

Pearson’s correlation coefficients between out-of-sample EEG data *X*_*t*_ and the average out-of-sample estimates 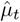 (obtained by averaging over individual realizations) were collected to measure the linear relationship between the two variables, with possible values ranging from -1 to 1. The Pearson’s correlation coefficient, defined below, states that values of -1 indicate strong negative linear correlation, whereas values of 1 indicate strong positive linear correlation. A Pearson’s correlation coefficient value of 0 indicates no linear correlation between two variables.

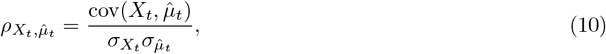

where 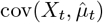 represents the covariance of *X*_*t*_ and 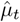, and *σ*_*i*_ represents the standard deviation of variable *i*.

Power spectra of individual realizations and out-of-sample EEG data were produced to analyze and compare corresponding frequency content across model types. Power spectral density was estimated using Welch’s method (`pwelch` in MATLAB 2024b) over a frequency range of 0 to 60 Hz with intervals of 0.1 Hz and sampling frequency of 256 Hz. The mean of each individual realization was removed from the signal before computing the power spectrum to avoid large spikes and focus solely on frequency composition.

Similarly, the mean of the EEG data in the out-of-sample period was removed beforehand. The mean power spectrum 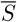 was obtained by averaging over individual realization power spectra and compared against the EEG data power spectrum *S* by using the mean squared error:

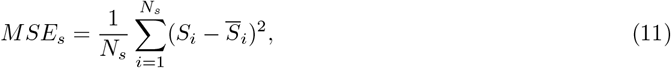

where *N*_*s*_ = 6001 is the number of observations in the frequency domain.

### Statistical Significance: Wilcoxon Rank-Sum Test

The mean squared error of the point-wise histograms, power spectrum errors and the Pearson’s correlation coefficients of all average out-of-sample estimates within each model type were aggregated across 15 participants, 4 trials, and 5 intervals to compare model performance. Predictions with lower mean-squared errors and large positive Pearson’s are favorable, indicating their corresponding models have an improved ability to adequately describe the EEG data and predict future values over other models. A pairwise Wilcoxon Rank-Sum test (**WRST**) was used to test the null hypothesis that the two sets of numbers were generated from the same probability distribution. Four model types (AR, ARX, ARMAX, SARMAX) indicate that six pairwise tests are necessary to compare every model type.

*WRST Statistical Testing Assumptions:*

- Mean squared errors and correlation coefficients between any two given model types are independent of each other.
- Mean squared errors and correlation coefficients are ordinal.
- Distributions of mean squared errors and correlation coefficients within a given model type across participants and trials are not necessarily normal.

WRST results were obtained in MATLAB 2024b using the `ranksum` function with an appropriate significance level *α*. Resulting *p*−values correspond to the probability of correctly rejecting the null hypothesis, but effect sizes further quantify the strength of any differences. Given two groups of sizes *n*_1_ and *n*_2_ (here *n*_1_ = *n*_2_ = 300), the WRST effect sizes were defined as the following, (Cohen, 2013; Tomczak and Tomczak, 2014):

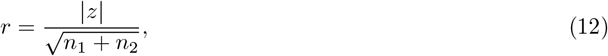

where *z* represents the *z*-score of the *U* -statistic: *z* = (*U − µ*_*U*_)*/σ*_*U*_ . Effect sizes were assigned a qualitative measure according to their magnitude by (Cohen, 2013; Tomczak and Tomczak, 2014), such that sizes within the range (0,0.2] were considered small, (0.2,0.5] were medium, and (0.5,0.8] or above were large.

Power spectra of out-of-sample estimations across all participants, trials, and intervals were subdivided into *δ* (1-4 Hz), *θ* (4-8 Hz), *α* (8-13 Hz), and *β* (13-30 Hz) frequency bands to determine model selectivity for certain frequency ranges and frequency selectivity for certain model type. A similar pairwise WRST testing scheme was used to determine if distributions within frequency bands and across model types are significantly different and if distributions across frequency bands and within model types are significantly different.

## Acknowledgments

We thank the VCU Breakthroughs Fund 2022–2024 for support (RRF, CMD, DJK, CL).

## Data Availability Statement

The EEG data are available upon request to the Krusienski Lab.

The code to implement the models and generate figures in this paper are freely available at https://github.com/RiccaRomano/TimeSeriesEEG.

## Appendix A

### Detailed In-Sample Fits

This section shows the results of in-sample fits for each of the models, as well as the corresponding mean-squared error for all 300 intervals (panel **A** in Figs A1–A3). Showing the complete details of all model terms for all 300 intervals; note that the ARMAX model in-sample fits are in the main text (Fig. 3).

### Detailed Out-of-sample Estimations

This section shows the out-of-sample estimation results (histogram errors, Pearson’s correlation, power spectrum errors) for each of the 4 models. The results of the in-sample fit are compared to the immediate period (3.5 s) out-of-sample (red model fit is cartoon for exposition, panel **A**). In each of the figures (A4 – A7), we show a comparison of time series values (point-wise), taking histograms of the out-of-sample period, showing the best, average, and worst fit (red) to data (black) out of all 300 tests (**B**). The right column shows the entire set of errors. The Pearson’s correlation of the out-of-sample estimates and data is also shown in each model following the same set-up (**C**). Since the frequency content of the signal is a key entity, we show a breakdown by the two m-sequences, showing the best and worst errors over all 5 intervals in a trial (**Di, Dii**).

As expected from the results in Figure 5, there are very few differences between the models, specifically between AR, ARX and ARMAX. The out-of-sample SARMAX estimates (Fig. A7) can be really off for some outliers that at times will not lead to statistically significant differences (exceptions are for the power spectra).

For SARMAX, the histograms in panel B can have very large mean-squared errors. Although indeed the Pearson’s correlation can have larger values (≈0.4), possibly indicating better performance (Fig A7C), the corresponding mean-squared error of the histogram of point-wise values for the higher Pearson’s correlation is very large (compare both right-most figures in Fig A7B and C). This is a shortcoming of Pearson’s correlation, necessitating the use of multiple metrics to assess estimation performance. Notably, the out-of-sample power spectra can be very bad compared to the actual data; see Figure A7Di, green interval 5, and Dii green intervals 4 and 5, where the overall power of the out-of-sample estimation is incredibly large compared to the actual data. Overall, the out-of-sample SARMAX estimates can be very bad despite more complexity; unfortunately, we currently do not know when the estimation will be bad since there are no obvious signatures.

**Figure A1:**
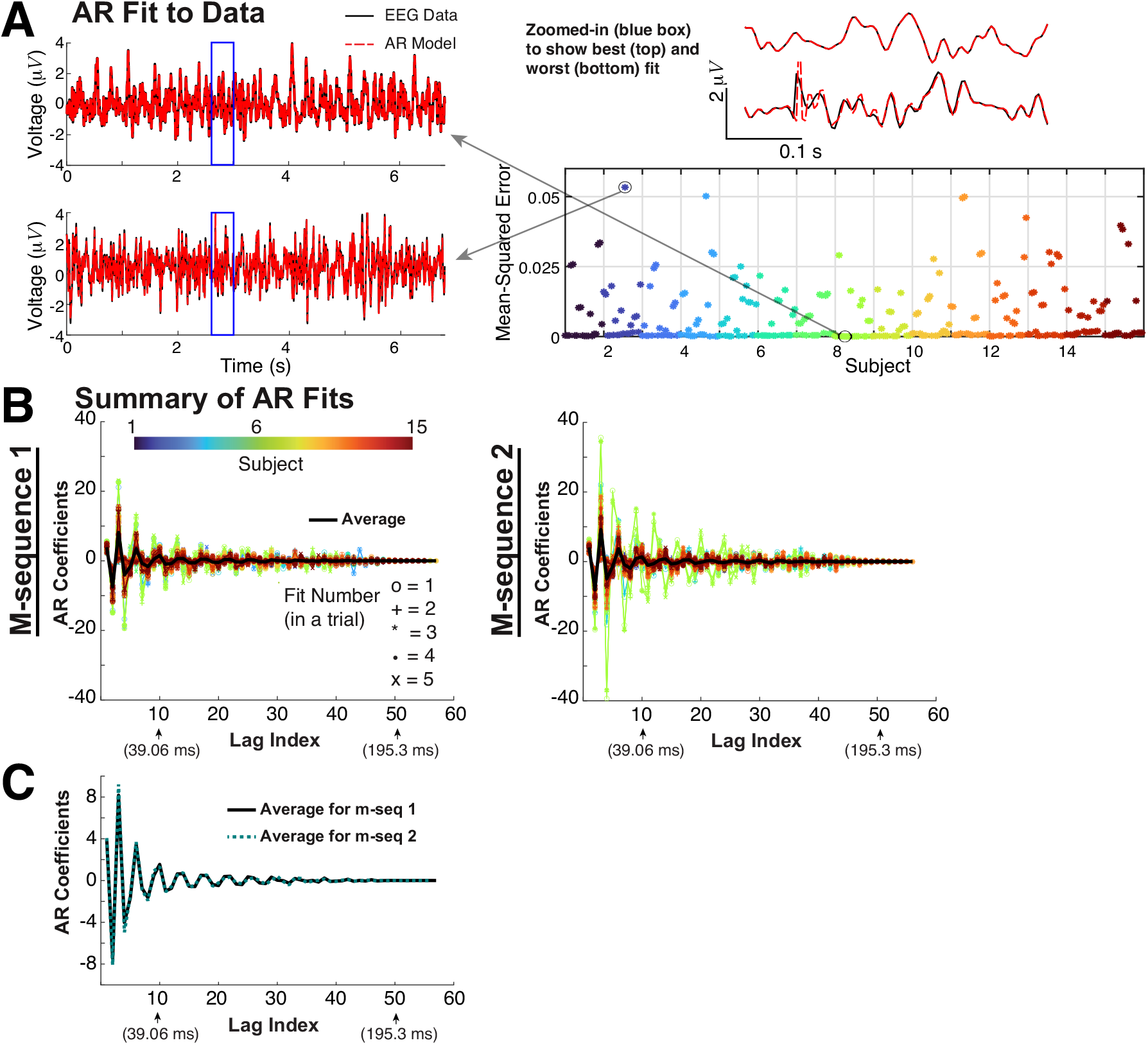
AR fit to data. **A**) Showing the best and worst in-sample fit among all 300 intervals. **B**) The AR terms for each of the 2 m-seqs; the black curve shows the average. **C**) The average AR values for each m-seq are similar (re-plotting black curves in B).

**Figure A2:**
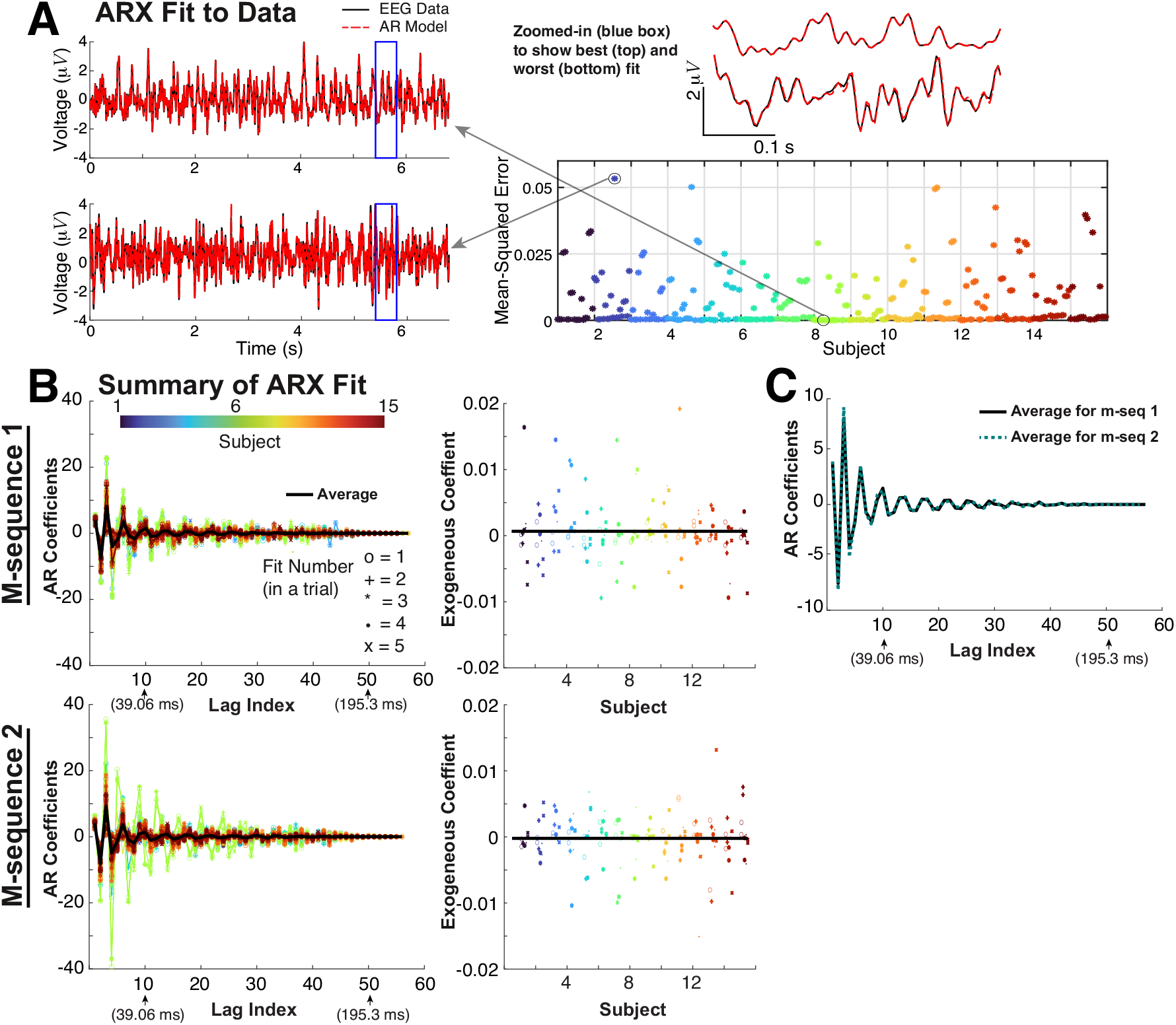
ARX fit to data. Same set-up as Fig A1 except the ARX model has an exogenous term. **A**) Showing the best and worst in-sample fit among all intervals. **B**) The AR terms for each of the 2 m-seqs; the black curve shows the average. Showing the exogenous values *η* and the average (black horizontal lines). **C**) The average AR values for each m-seq are similar.

**Figure A3:**
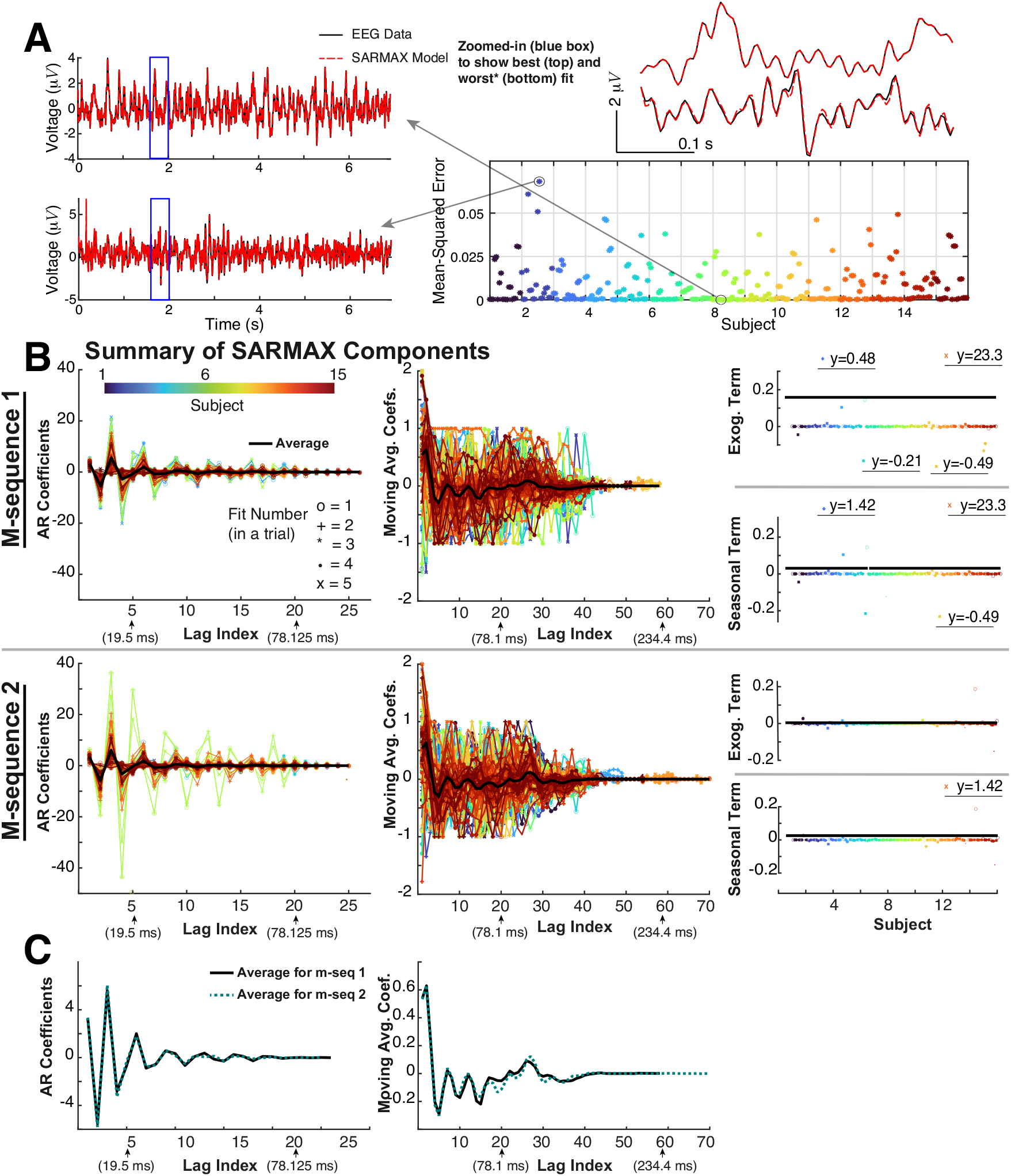
SARMAX fit to data. Same set-up as before but with additional model components. **A**) Showing the best and worst in-sample fit among all intervals. **B**) The AR terms for each of the 2 m-seqs; the black curve shows the average. Showing the exogenous values *η* and the seasonality coefficient, along with their average values (black horizontal lines). **C**) The average AR and MA values for both m-seqs are similar.

**Figure A4:**
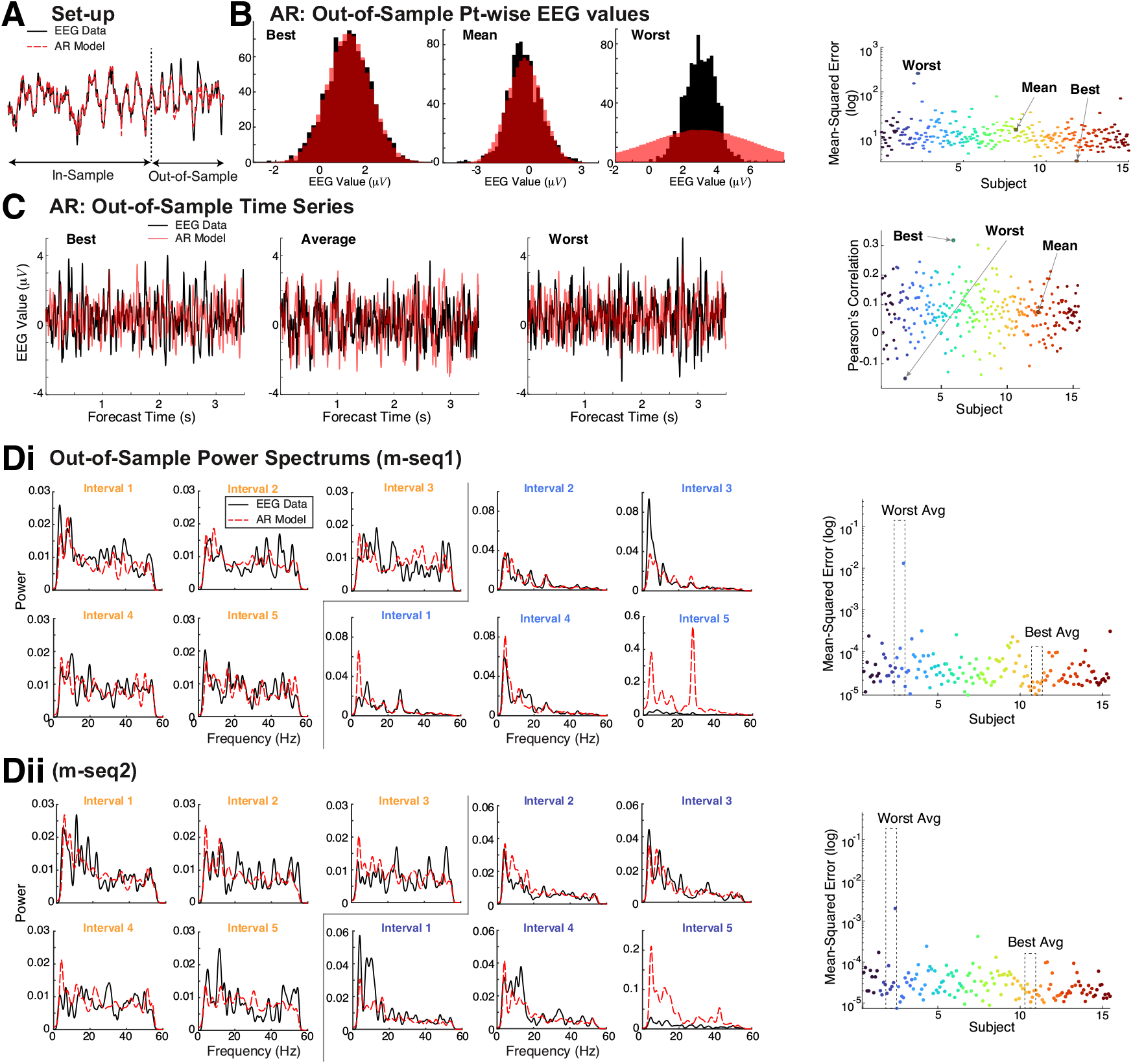
AR Out-of-sample Estimation Results. **A**) Set-up. **B**) Comparison of time series values via histograms of the best, average, and worst fit (red) to data (black) out of all 300. **C**) Comparison of EEG time series (black) with out-of-sample AR estimates (red) with the best, mean and worst Pearson’s correlation. The right panel shows all 300 Pearson’s correlation values. **Di**) With m-seq1, comparing the power spectrum of the data (black) and AR model fits (red-dashed). The right-most panel shows all 150 mean-squared error values (log-scale), and the 5 preceding columns show the best average mean-squared error within a trial (5 intervals of fitting and testing) with an orange heading and the worst average (dark blue heading). **Dii**) same as **Ai**) but for m-seq2. Right column: colors correspond to different subjects.

**Figure A5:**
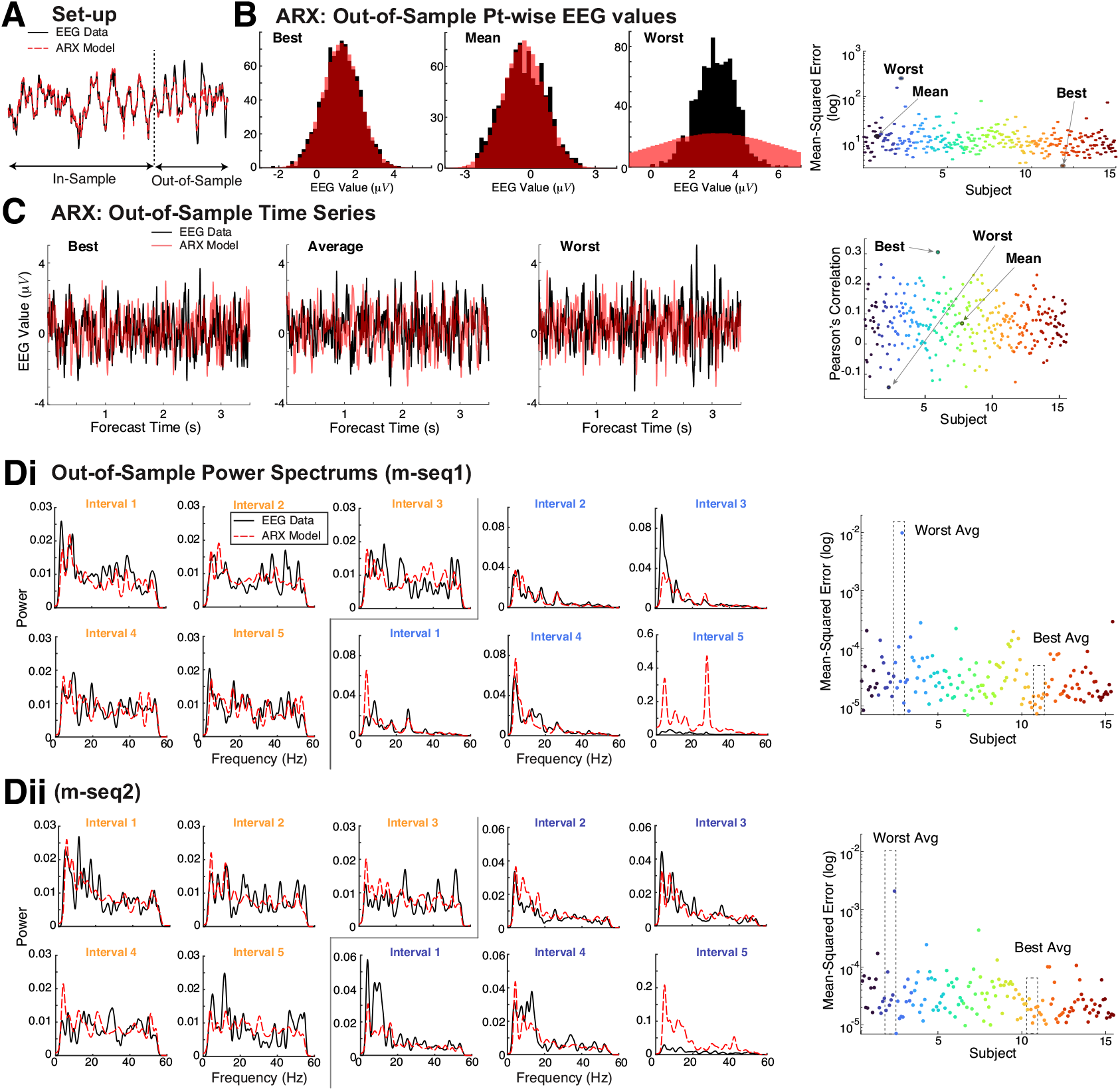
ARX Out-of-sample Estimation Results. **A**) Set-up. **B**) Comparison of time series values via histograms of the best, average, and worst fit (red) to data (black) out of all 300. **C**) Comparison of EEG time series (black) with out-of-sample ARX estimates (red) with the best, mean and worst Pearson’s correlation. The right panel shows all 300 Pearson’s correlation values. **Di**) With m-seq1, comparing the power spectrum of the data (black) and ARX model fits (red-dashed). The right-most panel shows all 150 mean-squared error values (log-scale), and the 5 preceding columns show the best average mean-squared error within a trial (5 intervals of fitting and testing) with an orange heading and the worst average (dark blue heading). **Dii**) same as **Ai**) but for m-seq2. Right column: colors correspond to different subjects.

**Figure A6:**
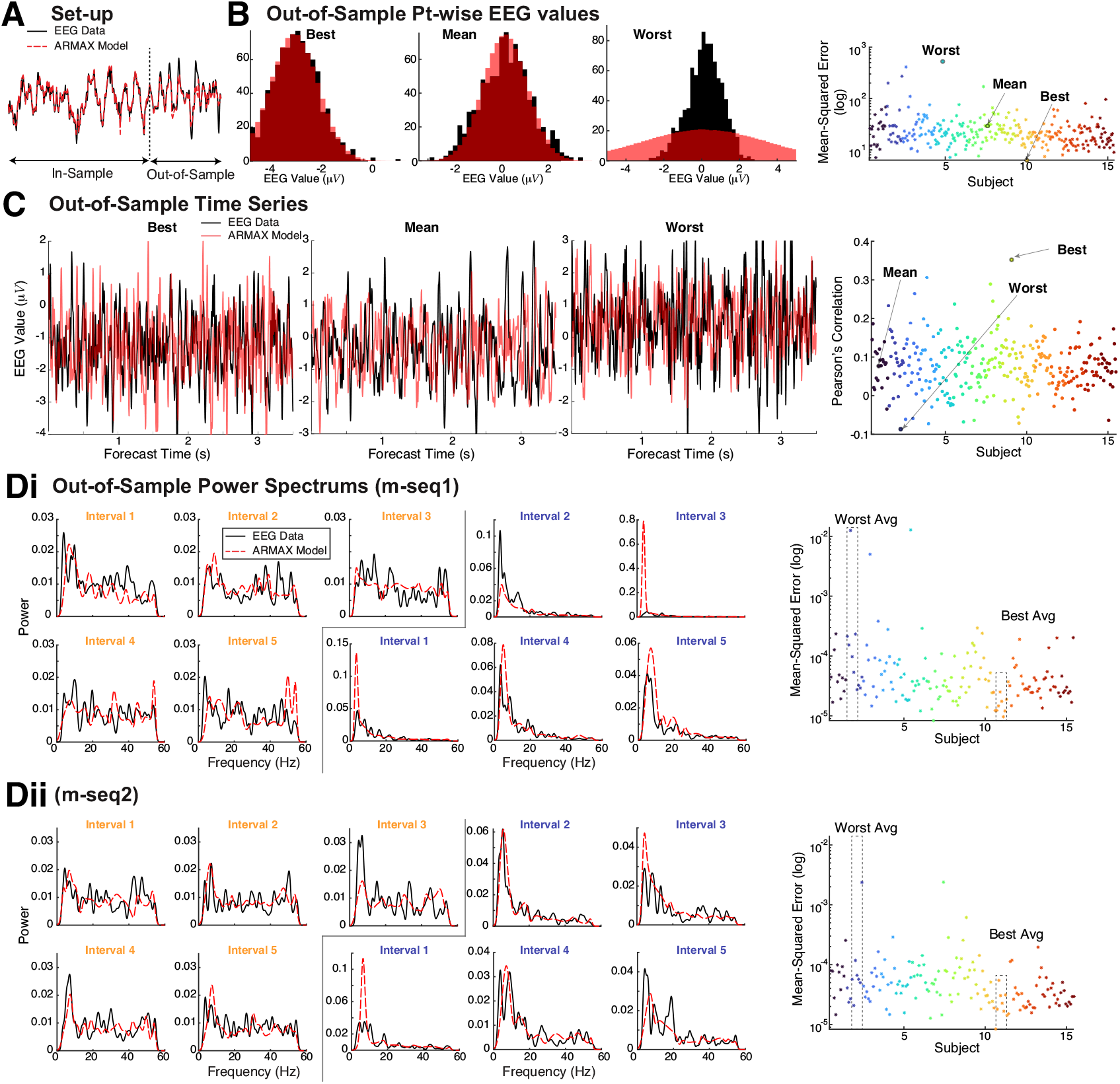
ARMAX Out-of-sample Estimation Results. **A**) Set-up. **B**) Comparison of time series values via histograms of the best, average, and worst fit (red) to data (black) out of all 300. **C**) Comparison of EEG time series (black) with out-of-sample ARMAX estimates (red) with the best, mean and worst Pearson’s correlation. The right panel shows all 300 Pearson’s correlation values. **Di**) With m-seq1, comparing the power spectrum of the data (black) and ARMAX model fits (red-dashed). The right-most panel shows all 150 mean-squared error values (log-scale), and the 5 preceding columns show the best average mean-squared error within a trial (5 intervals of fitting and testing) with an orange heading and the worst average (dark blue heading). **Dii**) same as **Ai**) but for m-seq2. Right column: colors correspond to different subjects.

**Figure A7:**
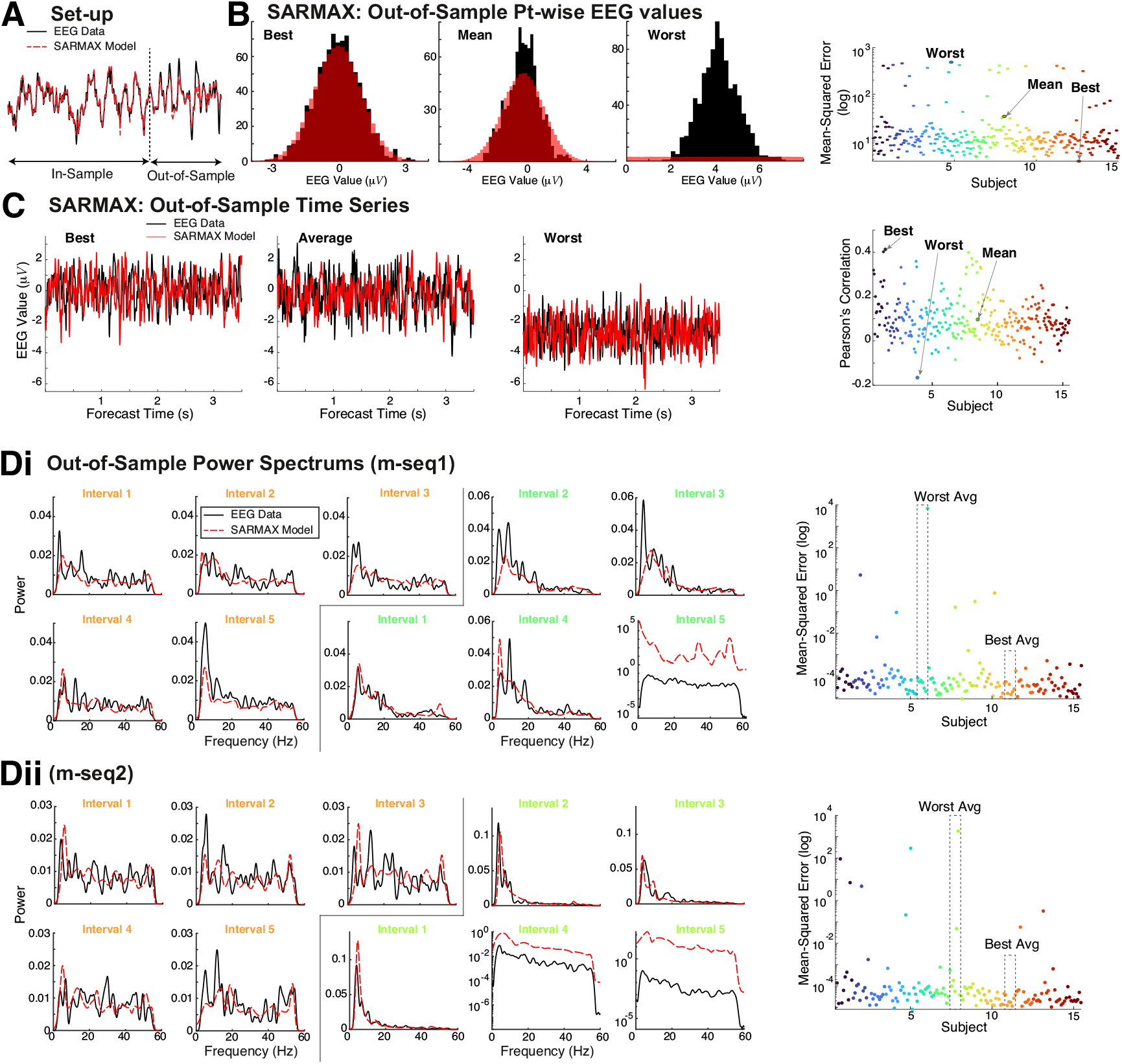
SARMAX Out-of-sample Estimation Results. **A**) Set-up. **B**) Comparison of time series values via histograms of the best, average, and worst fit (red) to data (black) out of all 300. **C**) Comparison of EEG time series (black) with out-of-sample SARMAX estimates (red) with the best, mean and worst Pearson’s correlation. The right panel shows all 300 Pearson’s correlation values. **Di**) With m-seq1, comparing the power spectrum of the data (black) and SARMAX model fits (red-dashed). The right-most panel shows all 150 mean-squared error values (log-scale), and the 5 preceding columns show the best average mean-squared error within a trial (5 intervals of fitting and testing) with an orange heading and the worst average (green heading). **Dii**) same as **Ai**) but for m-seq2. Right column: colors correspond to different subjects.

**Figure A8:**
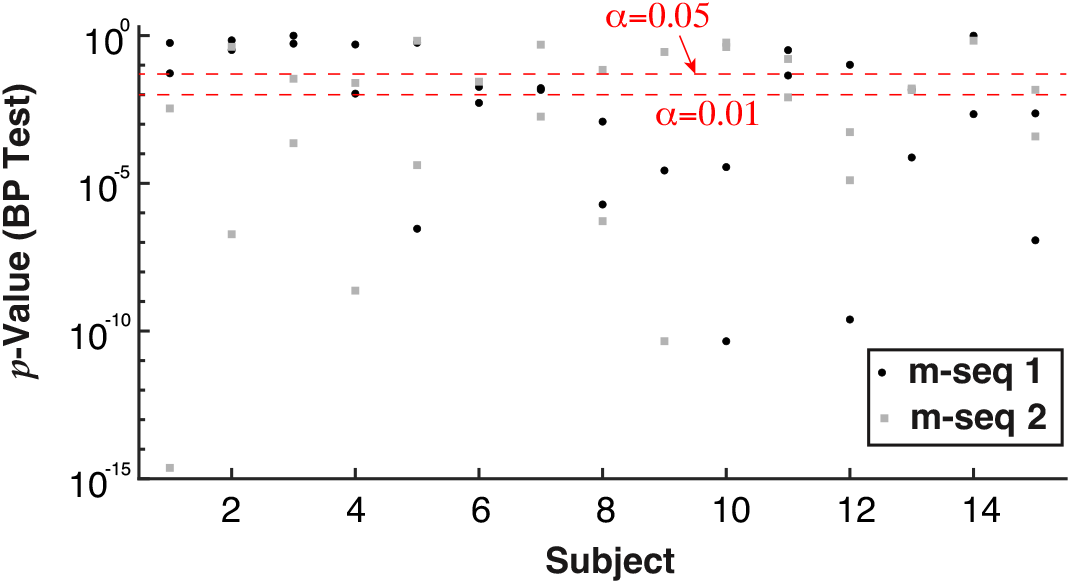
BP Test *p−*values. Whether a given time series is heteroscedastic (*p−*values above the significance level) or not does not depend on participant or m-sequence type (mseq-1 is a black circle, mseq-2 is a gray square).

